# Endosperm secretion remodels the maternal seed coat

**DOI:** 10.64898/2026.10.26.762064

**Authors:** Ke Yang, Wenbin Guo, Yuling Tang, Yue Li, Yunjun Zhu, Xiaoyuan Xi, Mingming Meng, Zeshuai Wang, Elin Axelsson, Hang Ye, Chunjiao Xia, Weibo Xie, Jialing Yao, Yang Zhu, Peitao Lü, Peng Zhao, Mengxiang Sun, Frédéric Berger, Ran Xu, Jing Li

**Affiliations:** School of Tropical Agriculture and Forestry, Sanya Institute of Breeding and Multiplication, Hainan University, Sanya 572025, China; College of Life Science and Technology, Huazhong Agricultural University, Wuhan 430070, China; Institute of Crop Science, Zhejiang University, Hangzhou 310058, China; Yazhouwan National Laboratory, Sanya 572024, China; Gregor Mendel Institute of Molecular Plant Biology, Austrian Academy of Sciences, Vienna BioCenter (VBC), 1030 Vienna, Austria; National Key Laboratory of Tropical Crop Breeding, Institute of Tropical Bioscience and Biotechnology and Sanya Research Institute, Chinese Academy of Tropical Agricultural Sciences, Sanya 572024, China; State Key Laboratory of Hybrid Rice, College of Life Sciences, Wuhan University, Wuhan 430070, China; Temasek Life Sciences Laboratory, National University of Singapore, Singapore 117604, Singapore

**Keywords:** *Arabidopsis thaliana*, endosperm, seed coat, apoplast, expansin, IKU–WRKY10, epigenetic regulation

## Abstract

In flowering plants and mammals, maternal tissues must accommodate the growth of the offspring they enclose. The maternal seed coat expands around the developing embryo and endosperm. Endosperm pressure and hormonal signals coordinate seed-coat development^1,2^, but whether the endosperm directly promotes maternal cell elongation remains unclear. Here we show that the coenocytic *Arabidopsis* endosperm directly supplies cell-wall-loosening proteins to maternal seed-coat walls. The seed-size-regulating HAIKU–WRKY10 pathway activates an endosperm secretory programme enriched in cell-wall-associated proteins. These proteins spread through a shared extracellular domain across multiple inner seed-coat layers, revealing an extensive functional maternal–filial interface through which the endosperm reaches maternal cells it does not directly contact. Among these proteins, three expansins are necessary and sufficient for maternal cell elongation and seed growth. The endosperm therefore promotes its own growth by enabling expansion of the maternal tissues that enclose it. Paternal DNA methylation and maternal Polycomb activity jointly regulate this programme, with Polycomb also required for timely shutdown of expansin expression. These findings link parental epigenetic states to maternal tissue expansion through proteins supplied by the offspring. This cross-generational control of growth parallels epigenetically regulated signaling from mammalian extraembryonic tissues^3^, suggesting a shared strategy for adapting the maternal environment to offspring development.

## Introduction

Embryos developing inside the maternal reproductive apparatus require both maternal nutrients and space to grow. In placental mammals, embryo-derived trophoblasts invade the decidua, remodel uterine vasculature and release signals that adapt maternal physiology to fetal demand^4,5^. Flowering plants face a similar physical constraint within the seed, where the embryo and endosperm develop inside the maternal seed coat. The endosperm is placed at the interface between the embryo and the maternal tissues. In *Arabidopsis*, the endosperm initially expands as a multinucleate coenocyte, a shared cytoplasm containing many nuclei that divide without cytokinesis, while the maternal seed coat cells elongate around it, together defining the space available to the embryo^6–8^.

Auxin produced by the endosperm after central-cell fertilization initiates seed-coat development from the ovule integuments^2^. It has been proposed that the pressure exerted by the rapidly expanding endosperm is sufficient to drive the subsequent growth of the maternal integument cells^1,9,10^. The endosperm and maternal integument cells lack direct cytoplasmic connections and are separated by a cutin-rich layer in the apoplast, the extracellular space that includes the cell walls^11^. Although this layer does not cover the entire interface, it has been considered a significant barrier to macromolecular movement between these tissues^11^. Whether the endosperm also supplies proteins that directly remodel maternal cell walls during seed growth has remained unclear.

Here we identify a secretory programme through which the endosperm promotes expansion of its maternal enclosure. We show that cell-wall-loosening expansins produced in the endosperm spread through a shared extracellular domain across multiple inner seed-coat layers and promote maternal cell elongation. Parental epigenetic states regulate this programme, linking gene regulation in the endosperm to growth of the surrounding maternal tissues.

## Results

### IKU–WRKY10 directs an endosperm secretory programme

The HAIKU (IKU) pathway active in endosperm provides a genetic link to investigate the relationship between endosperm growth and expansion of the maternal integuments. Mutations in *IKU1*, *IKU2* or *WRKY10/MINISEED3* (*MINI3*) restrict both endosperm growth and integument cell elongation^6,7,12,13^. To explore whether IKU promotes integument growth through extracellular factors acting across the endosperm–maternal interface, we used a public laser-capture atlas of developing seeds^14^ to identify genes co-expressed with *WRKY10*. Co-expressed genes were enriched for promoter-proximal W-box motifs (Fig. 1a,b and Extended Data Fig. 1a,b). The genes downregulated in *iku1*, *iku2* and *wrky10* showed highly significant overlap with one another and with the *WRKY10*-co-expressed set (Extended Data Fig. 1c). Intersecting these gene sets identified 70 IKU downstream genes (*IDGs*), defined at the ATH1 probe-set level (Fig. 1c, Extended Data Fig. 1c and Supplementary Tables 1–5).

**Figure 1.**
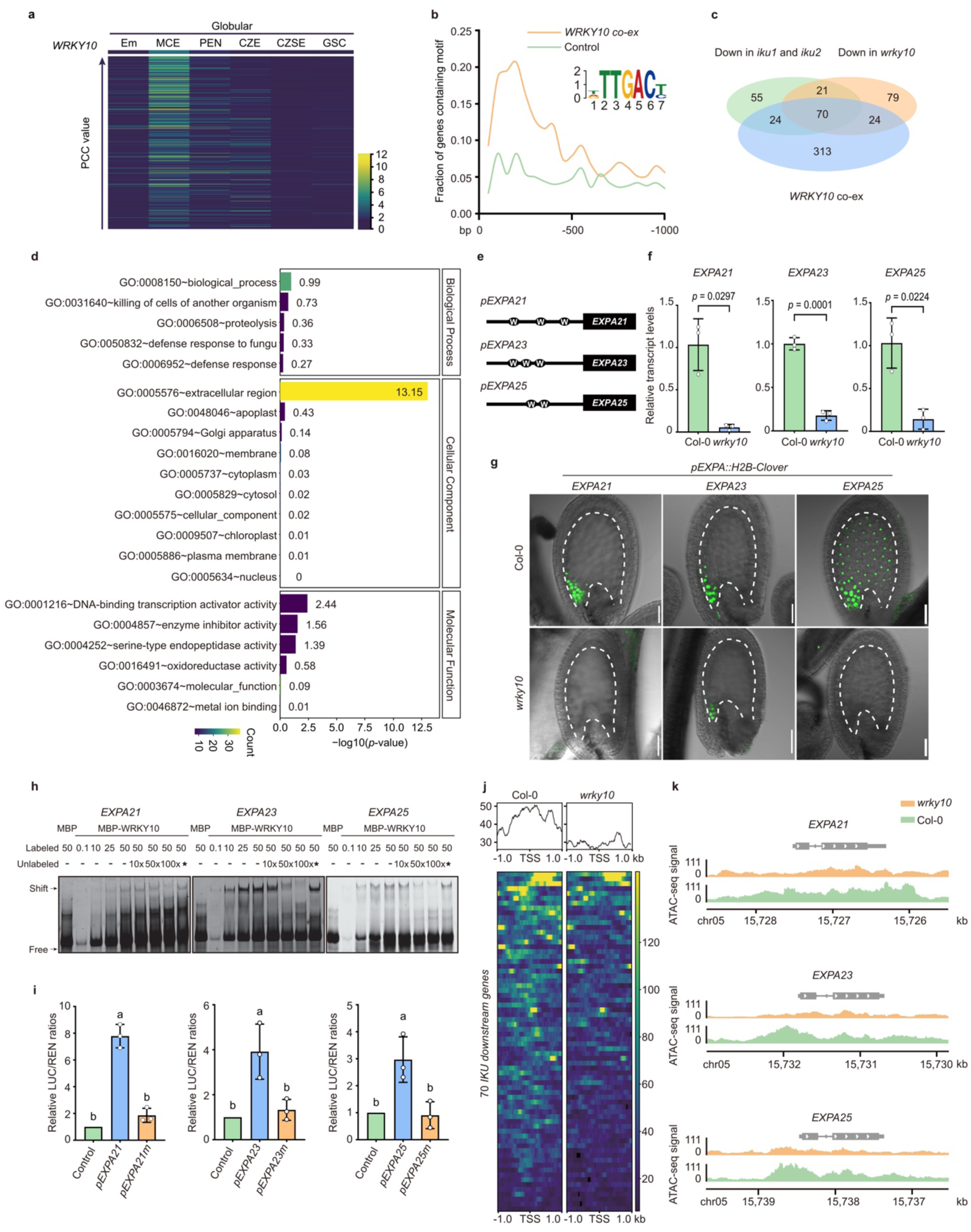
| The IKU–WRKY10 pathway activates an endosperm secretory programme. **a,** Expression of 431 ATH1 probe sets co-expressed with *WRKY10* in globular-stage seed compartments, ordered by decreasing Pearson correlation (PCC). Colour, normalized expression. Em, embryo; MCE, micropylar endosperm; PEN, peripheral endosperm; CZE, chalazal endosperm; CZSE, chalazal seed coat; GSC, general seed coat. **b,** W-box motifs within 1 kb upstream of *WRKY10*-co-expressed genes and 500 random controls. Inset, TTGACY motif. **c,** Overlap with probe sets downregulated in *iku1*, *iku2* and *wrky10*, defining 70 *IDGs*. **d,** Gene Ontology enrichment. Bar length, −log_10_(*P*); colour, *IDG* count. **e,** W-box positions within 500 bp upstream of the *EXPA21*, *EXPA23* and *EXPA25* start codons. **f,** RT–qPCR in 3-DAP Col-0 and *wrky10* seeds, normalized to *ACT11*. **g,** H2B–Clover reporters driven by the *EXPA21*, *EXPA23* and *EXPA25* promoters in Col-0 and *wrky10* seeds. **h,** MBP–WRKY10 binding to promoter fragments. Wild-type, but not W-box-mutant, competitors reduced binding. Numbers, molar excess. **i,** WRKY10-dependent activation of wild-type and W-box-mutant promoters in *Nicotiana benthamiana*. LUC was normalized to REN. **j,k,** ATAC-seq accessibility around *IDG* transcriptional start sites (±1 kb; **j**) and at the three *EXPA* loci (**k**). Data in **f,i**, mean ± s.d., three biological experiments. In **f**, exact *P* values are from two-tailed Student’s *t*-tests. In **i**, different letters indicate *P* < 0.05 by one-way ANOVA with Tukey’s test. Green, H2B–Clover; grey, transmitted light. Dashed lines, inner-integument boundary. Scale bars, 50 μm.

The proteins encoded by *IDGs* were strongly enriched for extracellular-region annotations (Fig. 1d). Strikingly, many *IDGs* encoded predicted cell-wall modifiers, including four expansins, five pectin modifiers and thirteen enzymes involved in cell wall-component metabolism (Supplementary Table 5). We focused on the cluster of *IDGs* encoding A-type expansins, which promote cell wall stress relaxation and cell expansion^15,16^ (Extended Data Fig. 2a,b). The expression of *EXPA21*, *EXPA23* and *EXPA25* increased after fertilization and peaked at approximately 3 days after pollination (DAP) (Extended Data Fig. 2c). Promoter reporters confirmed expression in the coenocytic endosperm. Activity was strongest in the micropylar region, decreased through the peripheral endosperm and was undetectable in the chalazal cyst (Extended Data Fig. 3).

The promoter of each gene contained W-box clusters that are known to bind WRKY transcription factors (Fig. 1e), and both transcript abundance and promoter-reporter activity were strongly reduced in *wrky10* seeds (Fig. 1f,g). We showed that WRKY10 bound W-box-containing promoter fragments in vitro, and that motif mutations impaired both binding and WRKY10-dependent activation in dual-luciferase assays (Fig. 1h,i and Extended Data Fig. 4a,b). Chromatin accessibility across the *IDG* set, including the expansin loci, was also reduced in *wrky10* endosperm nuclei (Fig. 1j,k). These results identify *EXPA21*, *EXPA23* and *EXPA25* as direct WRKY10 targets within a broader programme of cell-wall-associated genes expressed before the endosperm forms its own internal cell walls.

### Endosperm proteins reach maternal cell walls

The presence of a predicted N-terminal signal peptide in 38 of these proteins (Supplementary Table 5) suggested that they are exported and act outside the endosperm. To determine where endosperm-produced expansins accumulate, we expressed fluorescently tagged EXPA21, EXPA23 and EXPA25 under their native promoters. The expansins localized predominantly to maternal integument cell walls with comparatively weak fluorescence in the endosperm (Fig. 2a–f). More precisely, expansins occupied the cell walls of multiple inner-integument cell layers, including the parenchymatous ii1′ cells. In contrast, they were not detected in either the outer integument or the adjacent maternal tissues at the chalazal end (Fig. 2a,c,e). Apical views resolved a reticulate network of cell walls encircling most of the seed cavity (Fig. 2b,d,f). The fluorescence signal of these reporters was strongly attenuated in *wrky10* compared with wild type, consistent with their transcriptional dependence on WRKY10 (Extended Data Fig. 4c). Using the same approach, we detected fluorescently tagged proteins encoded by seven other *IDGs*, *SKS8*, *GH9B9*, *PMEI12*, *BG1*, *TBL15*, *PME39* and *EXPB4*, in maternal inner-integument cell walls (Extended Data Fig. 5). Thus, the IKU pathway controls the expression of a range of proteins that are deployed from the endosperm to surrounding cell walls, providing a potential link between endosperm growth and elongation of the maternal integument cells.

**Figure 2.**
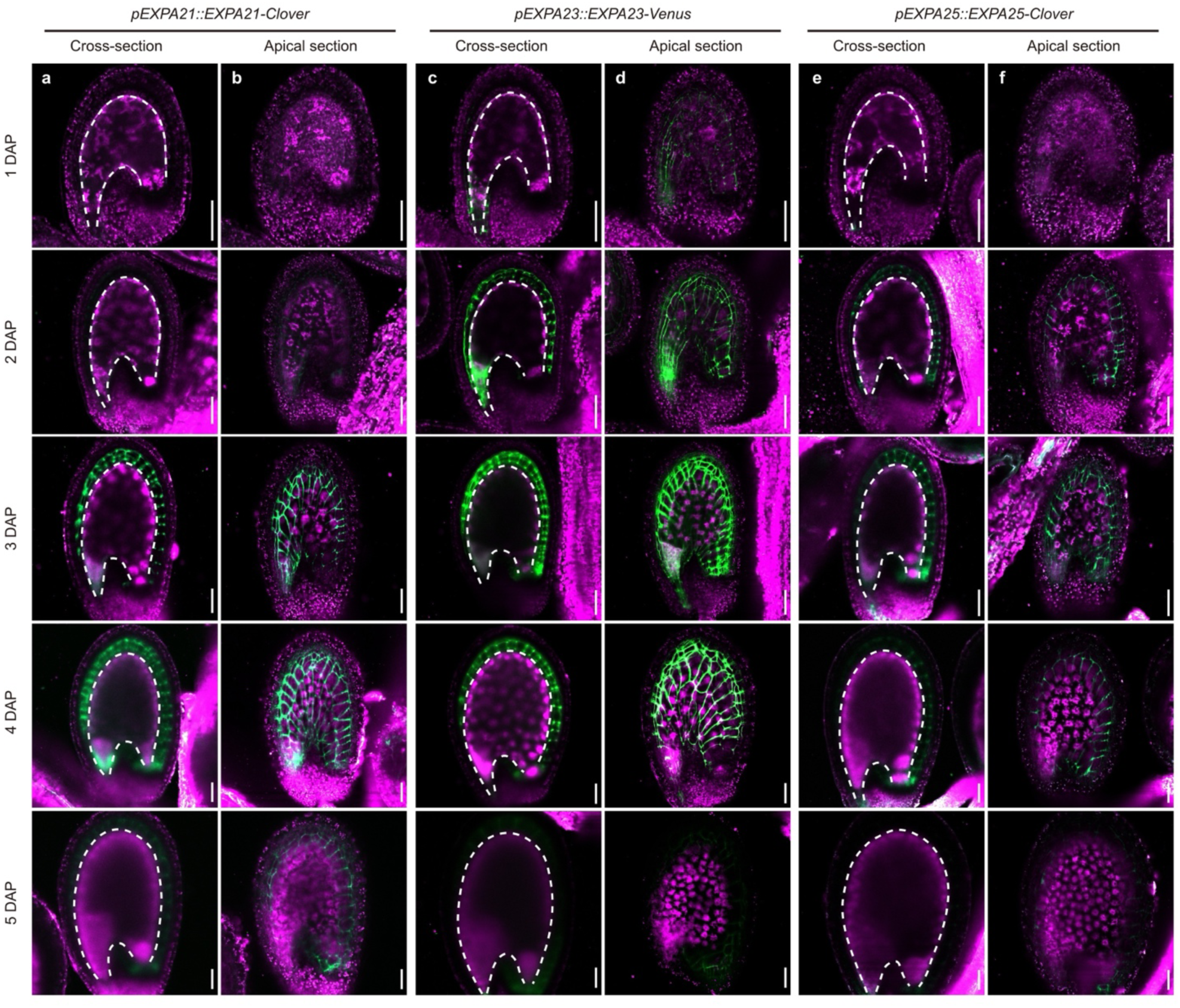
| Endosperm-derived expansins accumulate in maternal inner-integument cell walls. **a–f,** Confocal imaging of *Arabidopsis* seeds from 1 to 5 days after pollination (DAP) expressing native-promoter translational reporters for *EXPA21* (**a,b**), *EXPA23* (**c,d**) and *EXPA25* (**e,f**). EXPA21 and EXPA25 were fused to Clover and EXPA23 to Venus. Cross-sections (**a,c,e**) and apical sections (**b,d,f**) show the progressive appearance of fusion-protein signal in maternal inner-integument walls. Apical sections resolve a reticulate wall pattern surrounding most of the seed cavity. Green, EXPA–fluorescent-protein fusion signal; magenta, tissue autofluorescence. Dashed lines mark the endosperm-facing boundary of the maternal inner integuments. Scale bars, 50 μm.

To confirm that the proteins detected in maternal cell walls originated in the endosperm, we crossed *pEXPA21::EXPA21–Clover* males with females lacking the reporter. EXPA21–Clover still accumulated in maternal inner-integument cell walls (Fig. 3a), although expression of the reporter was confined to the endosperm (Fig. 3b). Similarly, paternally transmitted EXPA23 and EXPA25 reporters showed the same maternal-wall localization (Extended Data Fig. 6). We conclude that EXPA21, EXPA23 and EXPA25 are produced in the endosperm and move into maternal integument cell walls (Fig. 3c).

**Figure 3.**
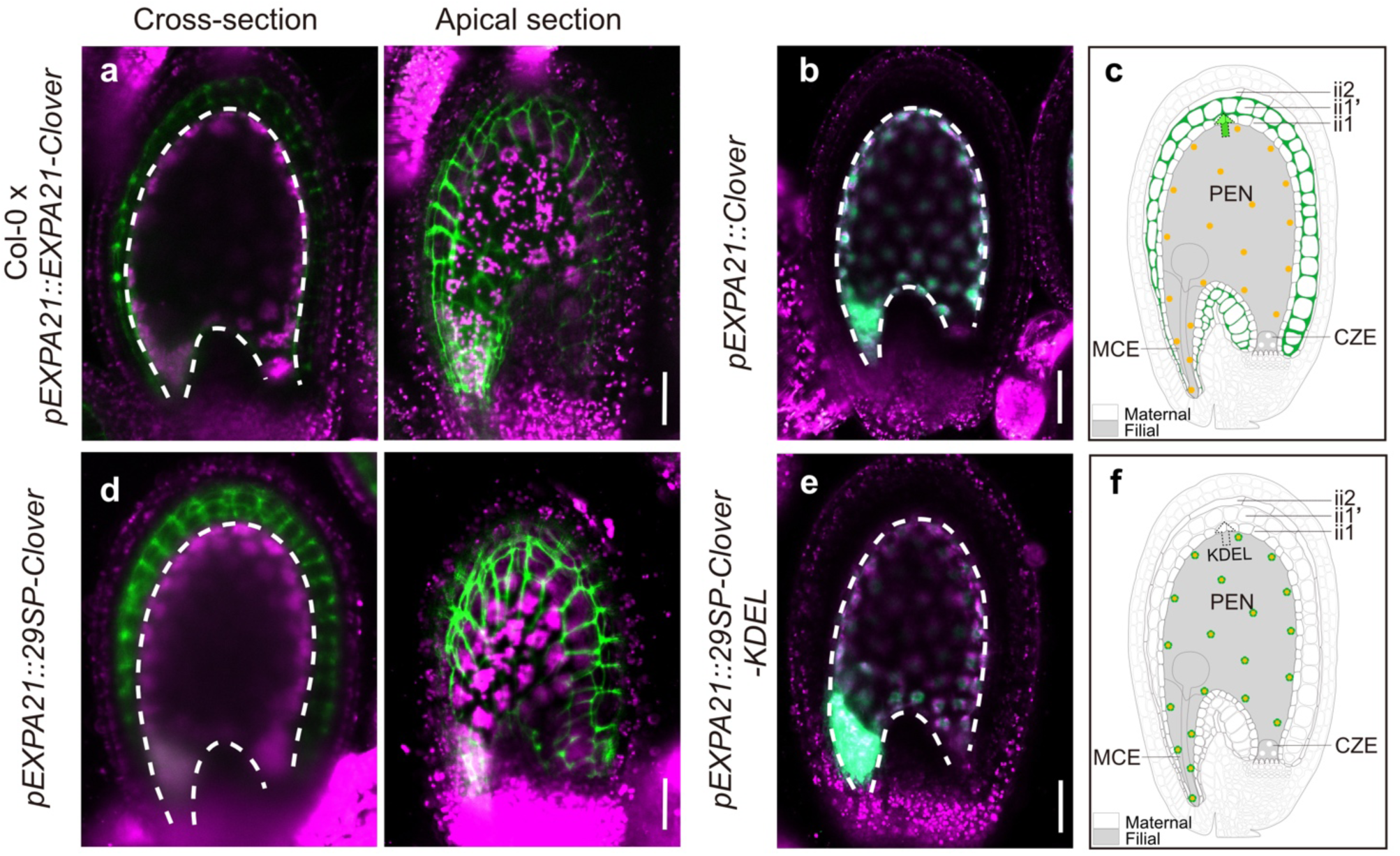
| Secretion delivers endosperm-derived proteins to maternal cell walls. **a,** Cross-sectional and apical views of seeds from Col-0 females pollinated by *pEXPA21::EXPA21–Clover* donors. **b,** *pEXPA21::Clover* reporter showing the expression domain of the *EXPA21* promoter. **c,** Schematic of **a,b**. **d,e,** The 29-amino-acid EXPA21 signal peptide directs Clover into maternal walls (**d**), whereas adding the endoplasmic-reticulum retention signal KDEL to 29SP–Clover confines fluorescence to the endosperm (**e**). **f,** Schematic of **d,e**. Orange nuclei denote expression source. Dark green denotes secreted-protein localization. Solid triangular arrows indicate direct adjacency of source and destination. Dashed arrows indicate observed redistribution through an unresolved route. Green, Clover; magenta, autofluorescence. Dashed lines mark the endosperm-facing inner-integument boundary. Scale bars, 50 μm. MCE, micropylar endosperm; PEN, peripheral endosperm; CZE, chalazal endosperm; ii1, innermost integument layer; ii1′ and ii2, subepidermal inner-integument layers.

### Endosperm secretion delivers proteins into maternal cell walls

We next asked whether this movement requires the conventional secretory pathway. The EXPA21 N-terminal signal peptide was sufficient to direct Clover to maternal inner-integument cell walls (Fig. 3d and Supplementary Table 5). Adding the endoplasmic-reticulum retention signal KDEL to 29SP–Clover confined fluorescence to the endosperm and prevented its localization to the maternal integuments (Fig. 3e). Deleting the signal peptide eliminated detectable fluorescence from the EXPA21 fusion driven by its native promoter. Even with the strong *FWA* promoter, which is active throughout the endosperm^17^, transcripts were detected but fusion-protein fluorescence remained undetectable (Extended Data Fig. 7). Conventional secretion is therefore necessary and sufficient to deliver these proteins from the endosperm into maternal inner-integument walls (Fig. 3f).

### Endosperm-derived expansins promote maternal growth non-cell-autonomously

Native expansins accumulated in the walls of the elongating inner integuments. To test whether these proteins promote maternal cell elongation, we generated an *expa21 expa23 expa25* triple mutant (*expa^tri^*) (Extended Data Fig. 8a,b). The mutant showed no visible vegetative defect under our conditions but produced smaller mature seeds resembling those of *wrky10* (Fig. 4a and Extended Data Fig. 8c–i). Combining *expa^tri^* with *wrky10* did not further reduce seed area (Fig. 4a). Together with direct transcriptional activation of these *EXPA* genes by WRKY10, this genetic non-additivity supports a role for the expansins downstream of WRKY10 in the IKU growth programme.

**Figure 4.**
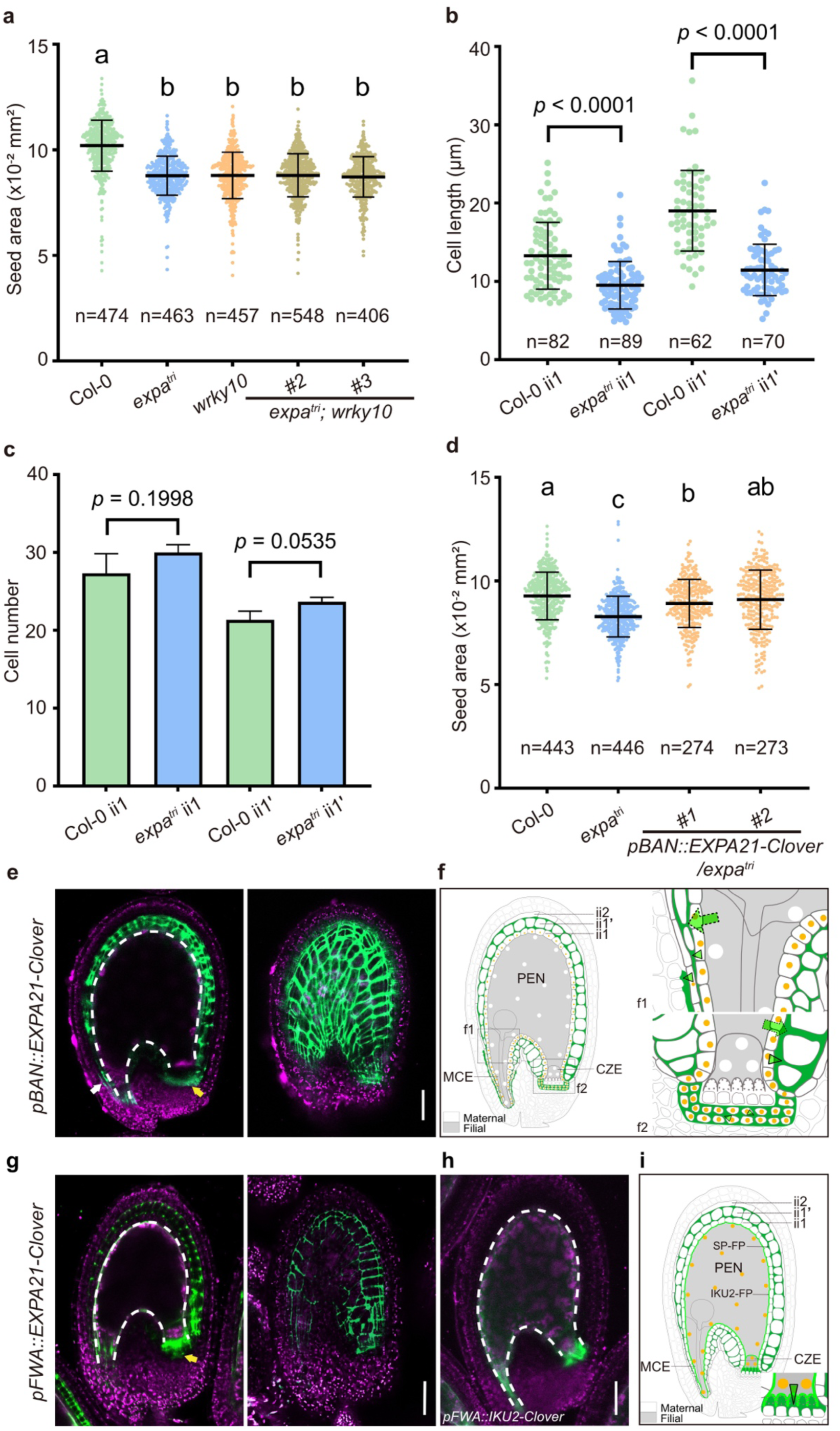
| Endosperm-derived expansins promote maternal growth non-cell-autonomously. **a,** Mature seed area of Col-0, *expa21 expa23 expa25* (*expa^tri^*), *wrky10* and combined mutants. **b,c,** Inner-integument cell length (**b**) and number (**c**) in Col-0 and *expa^tri^* at 3 DAP (ii1, ii1′). **d,** Seed area of Col-0, *expa^tri^* and two independent *pBAN::EXPA21–Clover* lines in *expa^tri^*. **e,** Cross-sectional and apical views of *pBAN::EXPA21–Clover* seeds. White and yellow arrowheads mark local micropylar ii2 and chalazal pigment-strand signal, respectively. **f,** Schematic of **e**, with micropylar (**f1**) and chalazal (**f2**) enlargements. The pigment strand joins the chalazal ends of ii1 and is separated from the endosperm cyst by nucellar tissue. **g,** Cross-sectional and apical views of *pFWA::EXPA21–Clover* seeds, showing inner-integument walls and an additional extracellular domain between the basal chalazal cyst and maternal nucellus. The yellow arrowhead marks this additional signal. **h,** Cross-sectional view of a *pFWA::IKU2–Clover* seed. The membrane-anchored reporter outlines the endosperm surface and elaborate basal cyst projections. **i,** Schematic of **g,h**, with an enlargement of the basal cyst–nucellus interface. Orange nuclei denote expression source. Dark green denotes secreted-protein localization, and bright green in **i** denotes membrane-localized IKU2. Solid triangular arrows indicate direct source–destination adjacency; dashed arrows, redistribution through an unresolved route. MCE, micropylar endosperm; PEN, peripheral endosperm; CZE, chalazal endosperm; ii1, innermost integument layer; ii1′ and ii2, subepidermal layers. In **a,d**, points are seeds from one representative plant per group, with concordant phenotypes in at least three independent plants; *n*, seed number; #, independent transgenic line. Points represent individual observations; summaries are mean ± s.d. In **a,d**, different letters indicate *P* < 0.05 by one-way ANOVA with Tukey’s test for the displayed seed distributions. Two-tailed Student’s *t*-tests were used in **b,c**, where exact *P* values are shown. In **e,g,h**, green indicates Clover signal, magenta indicates autofluorescence and dashed lines mark the endosperm-facing inner-integument boundary. Scale bars, 50 μm.

Consistently, *expa^tri^* seeds showed reduced seed coat cell elongation, similar to *iku* mutants^6,7^. Cells in both inner-integument regions examined, ii1 and ii1′, were shorter in *expa^tri^*than in wild type (Fig. 4b), whereas their number was unchanged (Fig. 4c). We then asked whether expressing *EXPA21* in the maternal integuments directly could rescue the small-seed phenotype of *expa^tri^*. We used the *BANYULS* (*BAN*) promoter, which is active mainly in ii1, the innermost integument layer of the maternal seed coat (Extended Data Fig. 9a)^18,19^. Expression of *pBAN::EXPA21–Clover* in *expa^tri^* restored seed size to values closer to wild-type seeds despite the absence of *EXPA21* expression in the endosperm (Fig. 4d). Expression of *EXPA21* in the maternal seed coat therefore compensated for the loss of endosperm-derived expansins. Consistent with this, EXPA21–Clover expressed in wild type under the *BAN* promoter recapitulated the broad inner-integument pattern of the native EXPA reporters (Fig. 4e,f and Extended Data Fig. 9). These results collectively establish that endosperm-derived expansins act non-cell-autonomously to promote maternal integument elongation and seed growth.

### The apoplast around the endosperm forms distinct interfaces with maternal tissues

Notably, the *pBAN::H2B–Clover* reporter confirmed *BAN* activity not only in the ii1 layer but also in a few micropylar ii2 cells and within the chalazal pigment strand (Extended Data Fig. 9a)^18,19^. These *BAN*-expressing cells form a tannin-producing maternal enclosure around the seed cavity, with the pigment strand joining the chalazal ends of ii1^18^ (Fig. 4f). EXPA21–Clover fluorescence remained local around the micropylar ii2 cells and the chalazal pigment strand. The pigment-strand signal nevertheless completed the fluorescent outline at the chalazal pole, where native EXPA reporters left a gap (Fig. 4e,f and Extended Data Fig. 9b).

Unlike ii1, which adjoins the main body of the endosperm, the pigment strand is separated from the cyst by nucellar tissue^18^. The proximal nucellar cells facing the pigment strand persist during seed development^20^. EXPA21–Clover secreted from the pigment strand did not reach the cyst-facing space (Fig. 4e,f), indicating an apoplastic barrier between the pigment strand and the cyst– nucellus interface. We therefore asked whether secretion from the cyst itself could reveal the apoplastic space on the endosperm side of this barrier.

To test this, we used the *FWA* promoter to extend EXPA21–Clover expression to the cyst, which naturally expresses many secreted-peptide genes^21^. This reproduced the broad inner-integument pattern and revealed additional extracellular signal directly between the basal cyst surface and the maternal nucellus (Fig. 4g). Membrane-anchored IKU2–Clover expressed from the same promoter outlined the elaborate cyst surface but did not spread through maternal cell walls (Fig. 4h). Basal cyst projections form a transfer-cell-like wall labyrinth at this well-characterized exchange interface^22–24^. Secretion from the pigment strand and cyst thus labelled opposite sides of the intervening nucellus. Together, these patterns reveal a compartmentalized maternal–filial apoplast, with an extensive inner-integument domain and a localized domain directly adjoining the cyst (Fig. 4i).

### Parental epigenetic states regulate the programme promoting maternal growth

The endosperm secretory programme enables the offspring to promote expansion of the maternal tissues that enclose it. Endosperm development is subject to parental epigenetic regulation involving DNA methylation and the Polycomb repressive complex 2 (PRC2)^25–27^. We asked whether these controls also regulate the secretory output that promotes maternal growth. Paternal DNA hypomethylation, which restricts seed growth^25^, significantly reduced expression of 62 out of the 70 *IDGs* in 3-DAP seeds resulting from crosses between hypomethylated *Met1cos* males and wild-type females (Fig. 5a and Supplementary Table 6). By contrast, maternal loss of *FERTILIZATION INDEPENDENT SEED 2* (*FIS2*), a PRC2 subunit, perturbed expression of 39 *IDGs*, with 19 upregulated and 20 downregulated (Fig. 5a and Supplementary Table 6). These responses suggest that paternal DNA methylation supports broad *IDG* expression, whereas maternal PRC2 activity regulates the balance of expression within the programme. When the two perturbations were combined, the *IDG* expression pattern shifted towards that of wild type, revealing a non-additive response (Fig. 5a, Extended Data Fig. 10 and Supplementary Table 6).

**Figure 5.**
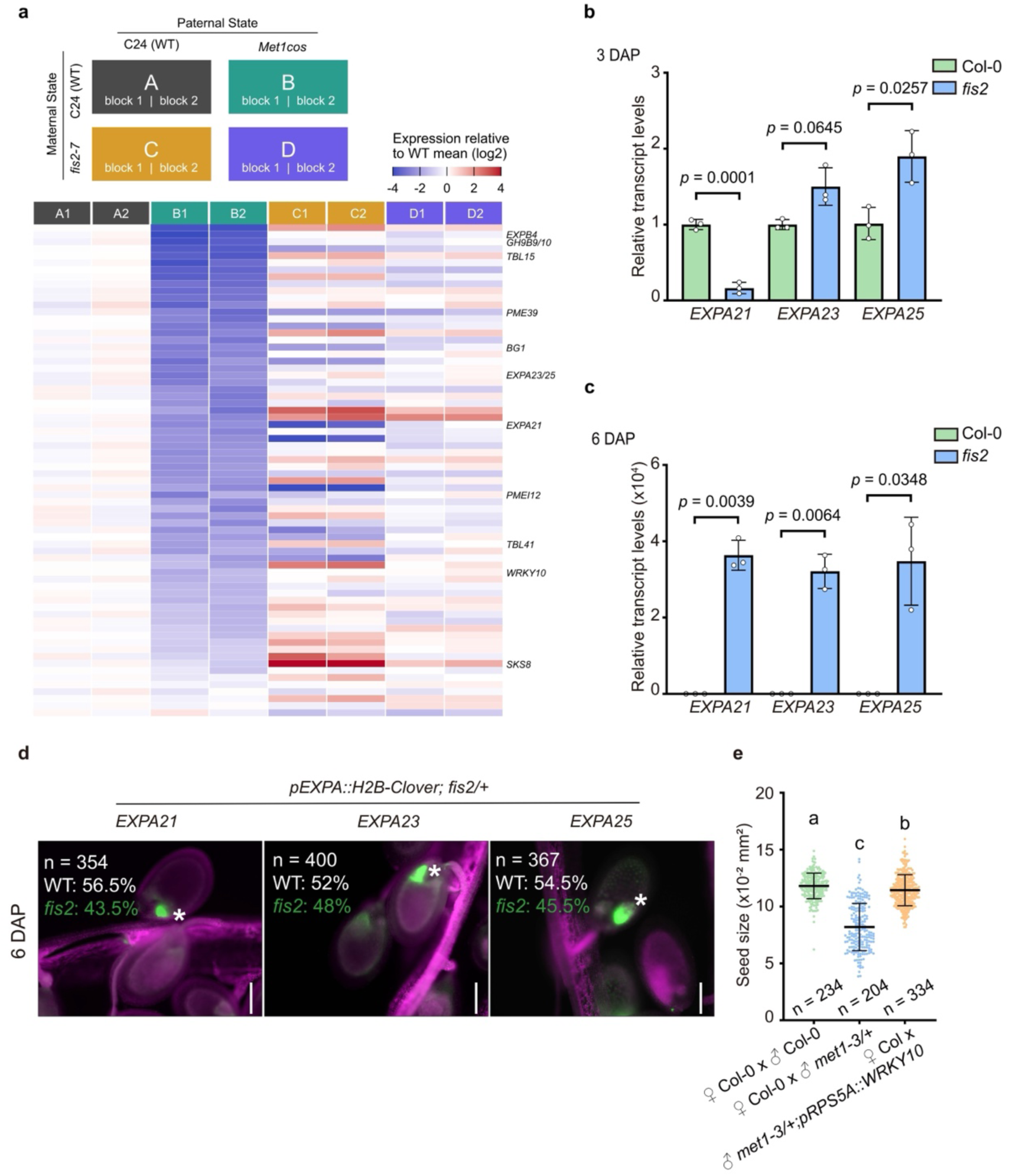
| Parental epigenetic states regulate the endosperm secretory programme. **a,** Expression of 70 *IDG*-associated ATH1 probe sets in 3-DAP seeds from a 2 × 2 design varying maternal *FIS2* and paternal methylation status. A, C24 × C24; B, C24 × *Met1cos*; C, *fis2-7* × C24; D, *fis2-7* × *Met1cos*. The female parent is listed first. Two biological sampling blocks were analysed per cross. Values are log_2_ expression relative to the mean of A. RMA-normalized data were analysed by blocked factorial limma, with Benjamini– Hochberg false-discovery-rate adjustment across ATH1 probe sets. **b,c,** RT–qPCR of *EXPA21*, *EXPA23* and *EXPA25* in Col-0 and *fis2* seeds (Col-0 background) at 3 DAP (**b**, normalized to *ACT11*) and 6 DAP (**c**). Data are mean ± s.d. of three biological replicates. Exact *P* values are from two-tailed Student’s *t*-tests. **d,** Representative 6-DAP seeds carrying *pEXPA21::H2B–Clover*, *pEXPA23::H2B–Clover* or *pEXPA25::H2B–Clover* in segregating *fis2/+* families (Col-0 background). Asterisks mark seeds retaining reporter activity. Seed numbers and genotype proportions are indicated in each panel. Green, H2B–Clover; magenta, tissue autofluorescence. Scale bars, 50 μm. **e,** Seed area from Col-0 females pollinated by Col-0, *met1-3/+* or *met1-3/+; pRPS5A::WRKY10* donors. Each point is one seed from one representative maternal plant (*n* = 234, 204 and 334 seeds, respectively). Horizontal lines indicate mean ± s.d. Phenotypes were concordant across at least three independent crosses. Different letters indicate *P* < 0.05 by one-way ANOVA with Tukey’s test for the displayed seed distributions. Seed area is expressed in 10^−2^ mm^2^.

FIS2 also regulated the developmental dynamics of expansin expression. At 3 DAP, *EXPA21* transcript abundance was reduced in *fis2*, whereas *EXPA23* and *EXPA25* were increased (Fig. 5b). At 6 DAP, all three transcripts were more abundant in *fis2* than in wild type (Fig. 5c). Their promoter reporters also remained active in *fis2* after activity had ceased in wild type (Fig. 5d). Loss of *FIS2* altered the balance of early expansin expression and prevented its timely shutdown. This temporal role is consistent with reported FIS–PRC2 regulation of the upstream *IKU2* and *WRKY10/MINI3* loci^28,29^.

Finally, we tested whether increased WRKY10 activity could counter the growth defect associated with paternal hypomethylation. A paternally transmitted *pRPS5A::WRKY10* transgene alleviated the small-seed phenotype caused by hypomethylated pollen (Fig. 5e). Together, these results connect parental epigenetic states to an endosperm secretory programme whose proteins promote expansion of the surrounding maternal tissues.

## Discussion

The endosperm actively makes room for its own growth by supplying proteins that loosen maternal cell walls. Before forming its own internal cell walls, the coenocytic endosperm deploys expansins through an IKU–WRKY10-dependent secretory programme. Fluorescent protein imaging and paternal genetic tracing establish that these proteins reach maternal cell walls deep within the seed coat, well beyond the endosperm-contacting cell layer. Together with loss-of-function and maternal complementation experiments, these findings demonstrate the non-cell-autonomous roles of endosperm-derived expansins in integument elongation and seed growth.

This secretory activity adds a biochemical component to mechanical models of seed growth. Creff and colleagues showed that endosperm pressure drives expansion but also induces tension-dependent reinforcement of the seed coat, particularly the inner-facing cell wall of the outer integument^1,9,10^. Expansins promote stress relaxation and irreversible cell wall extension under load^15,30^. Their delivery to the inner integuments gives the endosperm a means to influence how maternal cell walls respond to that pressure. Reduced expansin supply could help explain the restricted integument expansion of *iku2* seeds, which retain elevated endosperm pressure and undergo precocious seed-coat stiffening^1^. Thus, the endosperm provides both the force driving expansion and proteins that enable maternal cell walls to yield to that force.

The extracellular landscape revealed here also extends our understanding of the maternal–filial interface. Alongside the localized cyst–nucellus interface, endosperm secretion accesses an extensive apoplastic domain spanning multiple inner-integument layers around the main body of the expanding endosperm. The maternal–filial interface thus encompasses an extensive extracellular territory rather than being confined to the immediate contact between the two generations. Apoplastic barriers partition this landscape, defining which maternal cells are accessible from different sites of secretion. Beyond wall remodeling, this extensive domain provides a spatial framework for investigating nutrient and signal exchange. Peptide signaling already coordinates embryo and endosperm development^31,32^. Recognition of the embryo-derived peptide PIPL7 by IKU2 links endosperm patterning to embryonic development^32^, raising the possibility that the embryo also influences maternal growth through the endosperm secretory programme.

Although flowering plants and mammals diverged long ago, the endosperm and placenta have evolved convergent roles in shaping the maternal environment. The coenocytic organization of the developing endosperm also parallels the multinucleate organization of placental trophoblast syncytia. In mammals, the imprinted gene *Igf2* regulates placental hormone production and maternal metabolism^3^, and relaxin promotes remodeling of the maternal extracellular matrix^33^. In seeds, paternal DNA methylation and maternal Polycomb activity regulate the endosperm programme that supplies cell wall-loosening proteins to maternal tissues, with Polycomb also required for timely shutdown of expansin expression. Together, these parallels in cellular organization, offspring-driven modification of maternal tissues and parental epigenetic regulation suggest convergent strategies through which offspring modify their maternal environment to accommodate growth.

## Methods

### Plant material and growth conditions

The *Arabidopsis thaliana* accessions Columbia-0 (Col-0), Landsberg *erecta* (Ler) and C24 were used as wild-type backgrounds. The previously described *iku1* and *iku2* alleles were in Ler, and *wrky10/mini3* (SM_3_33099) was in Col-0^34^. The *fis2-7* (C24 background), *met1-3/+* and *Met1cos* materials were used in the parental epigenetic experiments described below. The *fis2* (Col-0), *expa21 expa23 expa25* triple mutant (*expa^tri^*) and *expa^tri^ wrky10* combinations were generated by CRISPR-Cas9 in Col-0.

Seeds were surface-sterilized in 75% ethanol for 5 min and 95% ethanol for 1 min, rinsed three times with sterile water, imbibed in sterile water for 24 h at 22 °C with gentle shaking and stratified at 4 °C for 2–3 d. Plants were grown in a 1:1:1 (v/v/v) mixture of Pindstrup peat moss (0–10 mm), commercial peat (3–5 mm) and vermiculite (2–4 mm), under 16-h light/8-h dark cycles, 120–150 μmol m^−2^ s^−1^ irradiance, approximately 22 °C and 60% relative humidity. *Nicotiana benthamiana* plants used for transient expression were grown under the same conditions.

For controlled crosses, stage-12 flower buds were emasculated^35^, allowed to mature for 48 h and manually pollinated. Days after pollination (DAP) refer to time after this controlled pollination.

### Bacterial strains

*Escherichia coli* DH5α was used for plasmid construction and BL21(DE3) for recombinant protein expression. Cells were cultured in LB medium with ampicillin (100 μg ml^−1^), spectinomycin (50 μg ml^−1^) or kanamycin (50 μg ml^−1^), as appropriate. *Agrobacterium tumefaciens* GV3101 was cultured at 28 °C in LB containing rifampicin (50 μg ml^−1^), gentamicin (25 μg ml^−1^) and the antibiotic required by the binary vector.

### Vector construction

Reporter constructs were generated by Gibson assembly^36^, and CRISPR constructs by Golden Gate assembly^37^. The kanamycin-resistance marker of the mini binary vector *pCB302*^38^ was replaced with spectinomycin resistance to generate *pSS100*. GFP or mCherry, driven by the mature-embryo-specific *At2S3* promoter, was inserted into the *pSS100* T-DNA to generate the seed-selection vectors *pSSG100* and *pSSR100*, respectively.

For transcriptional reporters, promoter fragments extending 1,906, 1,925 and 1,628 bp upstream of the ATG of *EXPA21*, *EXPA23* and *EXPA25*, respectively, were amplified from Col-0 genomic DNA and placed upstream of H2B–Clover in *pSSR100*. For native-promoter translational reporters, the same promoter fragments and corresponding genomic coding regions were fused in frame to Clover or Venus in *pSSR100* or *pSSG100*. Genomic promoter-plus-coding fragments were similarly used to generate translational reporters for *SKS8* (−1,508 to +2,707 bp), *GH9B9* (−1,603 to +1,767 bp), *PMEI12* (−2,070 to +522 bp), *BG1* (−2,072 to +1,780 bp), *TBL15* (−1,925 to +3,035 bp), *PME39* (−2,221 to +3,666 bp) and *EXPB4* (−2,037 to +1,660 bp), where coordinates are relative to the translational start site.

The *BAN* promoter (−1,906 to −1 bp) was used to drive H2B–Clover or EXPA21–Clover, and the *FWA* promoter (−3,274 to −1 bp) was used to drive EXPA21–Clover or IKU2–Clover. *EXPA21* and *IKU2* coding sequences for these constructs were amplified from Col-0 cDNA. The *RPS5A* promoter (−1,760 to −1 bp) was used to drive WRKY10–Clover.

For signal-peptide tests, the *EXPA21* promoter and coding sequence for the first 29 amino acids of EXPA21 were fused to Clover (*pEXPA21::29SP–Clover*). A promoter-only *pEXPA21::Clover* construct served as the no-signal-peptide control. The KDEL endoplasmic-reticulum retention sequence was appended to *pEXPA21::29SP–Clover* using the coding sequence AAGGACGAGCTG. EXPA21 signal-peptide-deletion constructs lacked the first 29 amino acids and were driven by either the *EXPA21* or *FWA* promoter.

The Cas9 cassette, including the *RPS5A* promoter, was based on *pKI1.1R*^39^. A guide-RNA scaffold amplified from *pHEE401*^40^ and a *ccdB* cassette were inserted into *pSSG100* to generate *pSSG_Cas9-ccdB*. Guide cassettes were amplified from *pCBC-DT1T2* and assembled as described previously^41^. The *fis2* (Col-0) mutant was generated using guides ATGGCTAGGAAGTCCATACG and AGATTCTTGTATACTGTAAG. The *expa21 expa23* double mutant was generated using guides TATGGTGATATCCATGGTGG and ATCGTCAAACACGGGTGCTT. The double mutant was subsequently transformed with two *EXPA25* guides, GACCCACAATGGTGTTTACC and AGGCTATGGTCTAGAGACAG, to generate *expa^tri^*, or with a *WRKY10* guide, ATGCTGGAGCAAACCGACAA, and an *EXPA25* guide, AGGCTATGGTCTAGAGACAG, to generate *expa^tri^ wrky10*. Guides were designed using CRISPR-GE^42^. All constructs and edited alleles were verified by Sanger sequencing. Primer sequences are provided in Supplementary Table 7.

### Stable transformation and line selection

Binary constructs were introduced into *A. tumefaciens* GV3101 by electroporation at 2.2 kV in a 0.2-cm cuvette. Arabidopsis plants were transformed by floral dip^43^, and transformants were identified using the GFP or mCherry seed-selection marker. More than 20 independent primary lines were screened for each reporter. At least three single-insertion lines with concordant localization patterns were evaluated, and one representative line is shown unless otherwise stated. Homozygous T3 or later generations were used. Reporter lines were crossed into mutant backgrounds and selected to homozygosity.

For genome editing, fluorescent T1 seeds were selected and target loci were sequenced. Non-fluorescent T2 seeds were selected to remove the Cas9 transgene, and Cas9-free homozygous lines were isolated in the T3 generation.

### Seed transcriptomes and co-expression

Spatial expression was analysed using the published laser-capture microdissection atlas of developing Arabidopsis seeds (GEO GSE12404)^14^. Figure 1a shows globular-stage seed compartments, whereas Extended Data Fig. 1a shows profiles from the pre-globular to mature stages. Expression values for embryo, micropylar endosperm, peripheral endosperm, chalazal endosperm, chalazal seed coat and general seed coat were used. Pearson correlation coefficients (PCCs) between *WRKY10* and the other represented ATH1 probe sets were calculated across the sampled seed compartments. The 431 probe sets with PCC ≥ 0.7, including *WRKY10*, were defined as the *WRKY10*-co-expressed set and ordered by decreasing PCC in the heatmap.

### Promoter motifs and *IDG* definition

Promoter sequences extending 1 kb upstream of the annotated transcriptional start site were scanned for W-box-containing motifs. Co-expressed genes were grouped by PCC interval, and frequencies of exact 5–8-bp motif matches without ambiguous bases were compared with other Arabidopsis promoters while accounting for local nucleotide composition and multiple testing (Fig. 1b and Extended Data Fig. 1b). The canonical W-box was represented as TTGACY, where Y denotes C or T^44^. Motif logos were generated with the MEME Suite^45^. For the focused promoter maps in Fig. 1e, W-box positions were plotted within 500 bp upstream of the translational start sites of *EXPA21*, *EXPA23* and *EXPA25*.

The *IDG* set was fixed before the parental epigenetic analysis. ATH1 probe sets downregulated in both *iku1* and *iku2* relative to Ler and in *wrky10* relative to Col-0 were intersected with the 431 *WRKY10*-co-expressed probe sets (Fig. 1c and Extended Data Fig. 1c). The 70 shared features were designated IKU downstream genes (*IDGs*; Supplementary Tables 1–5). Because *EXPA23* and *EXPA25* share an ATH1 probe set, the two genes are represented by a single microarray feature but were resolved by gene-specific RT–qPCR and reporter assays.

### Annotation and signal peptides

Gene Ontology annotations associated with extracellular-region and cell-wall-related functions were compiled for the fixed 70-*IDG* set using TAIR annotations (Fig. 1d). Protein functions were curated from TAIR and published annotations and grouped into the categories reported in Supplementary Table 5. N-terminal signal peptides were predicted with SignalP 6.0^46^. Signal-peptide sequence alignments were generated with MUSCLE5^47^.

### IKU discovery microarrays

Transcriptomes of 3-DAP Col-0, *wrky10*, Ler, *iku1* and *iku2* seeds were measured using Affymetrix ATH1 arrays. One biological sample per genotype was used for the Ler comparisons and two for the Col-0 comparison. Seeds were obtained by manual emasculation and controlled pollination, as described above. Pollinations were performed at the same time each day, and matched seed samples were harvested within a short time window. Each sample contained seeds dissected from at least 20 siliques. Total RNA was isolated using the PicoPure RNA Isolation Kit (Applied Biosystems, KIT0204), and array hybridization and scanning were performed as described previously^48^. Raw probe-level data were normalized by gcRMA. Differential expression was assessed by analysis of variance. Probe sets with *P* < 0.05 and fold change ≤0.5 were classified as downregulated, and probe sets with *P* < 0.05 and fold change ≥2 as upregulated. These data were used only to define the genotype-dependent sets contributing to Fig. 1c and Extended Data Fig. 1c; they were not combined with the RMA-normalized parental epigenetic experiment used in Fig. 5. Selected *IDGs* were validated by gene-specific RT–qPCR as described below.

### RNA isolation and RT–qPCR

Total RNA was extracted from ovules or seeds at the indicated DAP using the HiPure Plant RNA Mini Kit (Magen, R4151). First-strand cDNA was generated from approximately 500 ng RNA with the HiScript II 1st Strand cDNA Synthesis Kit with gDNA wiper (Vazyme, R212-01). Quantitative PCR was performed with 2× AceQ Universal SYBR qPCR Master Mix (Vazyme, Q511-02) on an Aria Real-Time PCR System (Agilent): 95 °C for 3 min, followed by 40 cycles of 95 °C for 5 s and 60 °C for 20 s. Transcript abundance was normalized to *ACT11* or *GAPC1* as specified in the figure legends. Each biological replicate was the mean of three technical reactions, and data are reported from three biological replicates. Primer sequences are listed in Supplementary Table 7.

### Electrophoretic mobility-shift assays

The *WRKY10* coding sequence (AT1G55600) was cloned into *pMAL-C2X* and expressed in *E. coli* BL21(DE3). Cultures were grown at 37 °C and 200 rpm to OD600 = 0.8, and expression of MBP or MBP–WRKY10 was induced with 0.5 mM isopropyl-β-D-thiogalactopyranoside for 16 h at 16 °C. Cells were lysed by low-temperature high-pressure homogenization (JN-10C, JNBIO) in 25 mM Tris-HCl pH 7.5, 300 mM NaCl, 5% glycerol and 0.5 mM dithiothreitol. Clarified lysates were passed through amylose resin (New England Biolabs), eluted with 10 mM maltose and concentrated using a 30-kDa-cut-off centrifugal device. Proteins were further purified by HiTrap Q ion-exchange chromatography with a 0–500 mM NaCl gradient in 25 mM Tris-HCl pH 8.0, followed by Superdex 200 Increase 10/300 GL size-exclusion chromatography in 25 mM Tris-HCl pH 8.0 and 2 mM dithiothreitol.

Electrophoretic mobility-shift assays were performed as described previously^49^ with minor modifications. Complementary oligonucleotides were annealed to generate 5′-FAM-labelled probes and unlabelled competitors. Binding reactions were incubated for 15 min at 22 °C in 25 mM Tris-HCl pH 8.0, 150 mM NaCl, 5 mM MgCl2, 1 mM dithiothreitol, 50 μg ml^−1^ salmon-sperm DNA and 10% glycerol. Complexes were resolved on 6% native polyacrylamide gels in 0.5× Tris-glycine buffer at 120 V for 1 h and imaged with an Amersham Typhoon scanner. Unlabelled competitors were used at 10-, 50-or 100-fold molar excess. Mutant competitors carried a TGAC-to-AGAC substitution.

### Dual-luciferase assays

Promoter fragments spanning −500 to −1 bp relative to the ATG of *EXPA21*, *EXPA23*, *EXPA25*, *IKU2* and *WRKY10* were inserted into *pGreenII 0800-LUC*^50^. W-box-mutant reporters contained a TGAC-to-AGAC substitution. *WRKY10* was expressed from the CaMV 35S promoter in *pFGC5941*^51^. Reporter and effector plasmids were introduced into *A. tumefaciens* GV3101(*pSoup*). Cultures at OD600 = 1.0 were washed twice and resuspended in 10 mM MES pH 5.7, 10 mM MgCl2 and 150 μM acetosyringone before co-infiltration into *N. benthamiana* leaves. Firefly and Renilla luciferase activities were measured 2 d later with the Dual-Luciferase Reporter Assay System (Promega, E1910) and an Infinite 200 PRO luminometer (Tecan). LUC/REN ratios were calculated for three independent infiltration experiments. Reporter and effector designs are shown in Extended Data Fig. 4a,b.

### ATAC-seq and analysis

ATAC-seq was performed once on 3-DAP Col-0 and *wrky10* seeds as described previously^52^ with modifications. Dissected seeds were chilled and chopped in 45 mM MgCl2, 30 mM sodium citrate, 20 mM MOPS pH 7.0 and 0.1% Triton X-100. Homogenates were passed through 30-μm mesh and stained with DAPI. After FSC-A/SSC-A gating, single nuclei were selected using DAPI-H versus DAPI-W, and the 6C population was gated on DAPI-A histograms using BD FACSDiva v8.0.1. Approximately 100,000 nuclei with 6C DNA content were isolated with a FACS Aria SORP flow cytometer (BD), pelleted at 500g for 5 min at 4 °C and tagmented with Tn5 transposase for 30 min at 37 °C. DNA was purified with the MiniBEST DNA Fragment Purification Kit v4.0 (TaKaRa, 9761), PCR-amplified and sequenced on an Illumina NovaSeq platform.

Reads were aligned to TAIR10 with Bowtie2 v2.5.4^53^. Properly paired alignments with mapping quality >30 were retained using SAMtools v1.21^54^, and PCR duplicates were removed with Picard Tools v3.3.0. Peaks were called with MACS2 v2.2.9.1 without model building, retaining one duplicate and generating SPMR-normalized bedGraph and summit outputs at *q* < 0.05^55^. Figure 1j displays accessibility within ±1 kb of the transcriptional start sites of the 70 *IDGs*; Fig. 1k shows genome-browser tracks at the three *EXPA* loci.

### Microscopy and protein-source tracing

Developing seeds were mounted directly and imaged with a Zeiss LSM 880 confocal microscope. Clover was excited at 488 nm and collected at 499–525 nm; Venus was excited at 514 nm and collected at 529–552 nm; RFP was excited at 543 nm and collected at 595–610 nm. Tissue autofluorescence was collected at 660–680 nm. Images were processed in ZEN v3.6 (Zeiss) using the same display settings for directly compared genotypes or constructs. Cross-sectional and apical optical views were assigned from seed anatomy. Whole seeds or representative anatomical regions were selected for imaging. Unless otherwise specified, images are from 3-DAP seeds. At least three independent transgenic lines were inspected for each reporter, with concordant localization patterns, and representative images are shown.

To trace a filial source independently of the maternal genotype, Col-0 pistils were pollinated with plants carrying native-promoter EXPA21, EXPA23 or EXPA25 translational reporters. Because the seed coat is entirely maternal, fluorescence from the paternally inherited transgene in maternal integument walls reports redistribution from the fertilization products. Secretion dependence was tested by comparing promoter-only Clover, EXPA signal-peptide–Clover, signal-peptide–Clover–KDEL and EXPA21 lacking its N-terminal signal peptide. The signal-peptide deletion was examined first under the native *EXPA21* promoter and then under the broader *FWA* promoter. Transcript accumulation from independent *pFWA::EXPA21^Δ29SP^–Clover* lines was assessed by cDNA amplification. The spatial domains in Fig. 4g–i were compared using endosperm-driven *pFWA::EXPA21–Clover* and membrane-anchored *pFWA::IKU2–Clover*. Maternal *pBAN::EXPA21–Clover* localization is shown in Fig. 4e,f. The diagrams in Fig. 3c,f and Fig. 4f,i summarize the adjacent microscopy panels. Orange nuclei denote the expression source, dark green denotes secreted-protein localization, and bright green in Fig. 4i denotes membrane-localized IKU2–Clover. Solid triangular arrows denote a source directly adjacent to the destination, and dashed arrows denote observed redistribution through an unresolved extracellular connection.

### Histology and integument measurements

Seeds at 3 DAP were fixed in FAA (50% ethanol, 5% acetic acid, 3.7% formaldehyde and 41.3% water) and vacuum infiltrated for 30 min. Samples were dehydrated through graded ethanol and xylene in a Leica TP1020 tissue processor, infiltrated with paraffin at 65 °C, embedded with a Leica HistoCore Arcadia system and sectioned at 4 μm on a Leica RM2265 microtome. Sections were floated at 40 °C, mounted, dried at 60 °C, dewaxed, rehydrated and stained with toluidine blue for 2–5 min. Inner-integument cell length and cell number in the ii1 and ii1′ regions were measured using automated image analysis in ImageJ^56^. The 3-DAP stage was selected because differences in inner-integument elongation are detectable in *iku* seeds at this stage^7^.

### Seed and vegetative phenotyping

For all mature-seed-area experiments in Fig. 4a,d and Fig. 5e, plants were grown together and sampled using the same procedure. Siliques were collected from the middle of the primary inflorescence after excluding the first and last five siliques; at least ten remaining siliques were collected from each plant when yellow-brown and beginning to dehisce. At least three morphologically comparable plants were examined for every genotype, transgenic line or controlled-cross condition. Siliques were dried at 37 °C for 7 d. Seeds were dispersed on white paper and imaged with a Nikon SMZ1000 stereomicroscope. Images were converted to 16-bit greyscale, thresholded and analysed with the ImageJ Analyze Particles function^56^. Incorrectly identified objects and unresolved clusters of overlapping seeds were removed after automated detection. Apart from these image-segmentation errors, no other experimental measurements were excluded. The current panels show all retained single-seed measurements from one representative plant for each group, and the direction of the phenotype was concordant among at least three independent plants. At least 200 seeds were analysed per group. A number sign denotes an independently derived transgenic line.

Whole plants were photographed 2 and 6 weeks after transplantation, and siliques were photographed at 7 DAP. Silique length was measured with digital callipers and seed number was counted manually in opened siliques. At least three independent plants and at least 20 siliques per plant were evaluated for vegetative and silique traits.

### Parental epigenetic microarrays

The parental epigenetic experiment in Fig. 5a and Extended Data Fig. 10 was analysed independently of the discovery microarrays above. Seeds were collected at 3 DAP in a balanced 2×2 crossing design with four conditions: A, C24 female × C24 male; B, C24 female × hypomethylated *Met1cos* male; C, *fis2-7* female × C24 male; and D, *fis2-7* female × *Met1cos* male. Two independent biological sampling blocks were analysed for every cross, giving eight ATH1 arrays: CC.CEL and CC2.CEL (A), CM.CEL and CM2.CEL (B), FC.CEL and FC2.CEL (C), and FM.CEL and FM2.CEL (D). All eight arrays passed the recorded array-image, RNA-degradation, relative-log-expression and normalized-unscaled-standard-error checks; no array was excluded.

Raw CEL files were processed in R 4.6.1 with affy v1.90.0. Probe-level values were background-corrected, quantile-normalized and summarized by robust multi-array average (RMA)^57,58^. ATH1 probe sets were annotated with ath1121501.db. Differential expression was analysed with limma v3.68.5^59^ using the blocked factorial model

“expression ∼ block + maternal FIS2 status + paternal MET1 status + maternal FIS2 status × paternal MET1 status”.

The pre-specified contrasts were B−A, C−A, D−A and the interaction D−C−B+A. Moderated statistics were obtained with empirical Bayes shrinkage, and false-discovery rates were calculated by Benjamini–Hochberg adjustment across all ATH1 probe sets within each contrast^60^. The fixed list of 70 *IDG*-associated probe sets was then extracted without re-selection. For Extended Data Fig. 10a, the additive prediction was (B−A) + (C−A) and the observed double-perturbation response was D−A. A probe set was designated closer to wild type when |D−A| < |(B−A) + (C−A)|. Extended Data Fig. 10b was restricted to the 35 probe sets for which both B−A and C−A were negative. The complete design matrix, contrasts, probe-level estimates and software session record accompany the analysis source data.

### FIS2-dependent temporal expression

Native *EXPA21*, *EXPA23* and *EXPA25* promoter reporters were crossed into the new *fis2* allele generated in Col-0 and imaged at 3 and 6 DAP using the confocal settings above. Wild-type and mutant seeds carrying the same reporter insertion were compared to assess persistence beyond the normal early expression window (Fig. 5d). Transcript abundance at 3 and 6 DAP was assessed by RT–qPCR in Col-0 and the same Col-0 *fis2* allele as described above (Fig. 5b,c).

### WRKY10 rescue crosses

For Fig. 5e, Col-0 maternal plants were pollinated with Col-0, *met1-3/+* or *met1-3/+*; *pRPS5A::WRKY10* pollen donors. The *met1-3/+* rescue experiment is genetically independent of the *Met1cos* material used for the transcriptome factorial design. Mature seed area was measured as above. This experiment tests whether increasing paternally transmitted WRKY10 activity shifts growth towards the control. It does not test allele-specific regulation of the endogenous locus.

### Phylogenetic analysis

The amino-acid sequences of 35 annotated Arabidopsis expansins were obtained from TAIR10; *EXPA19*, annotated as a pseudogene, was excluded. Sequences were aligned with ClustalW^61^, and a maximum-likelihood tree was inferred in MEGA X^62^ with 1,000 bootstrap replicates.

### Statistics and reproducibility

Statistical tests for molecular, cellular and seed phenotypes were performed in GraphPad Prism v8.0.0 unless otherwise stated. Two-group comparisons used two-tailed Student’s *t*-tests. Comparisons among three or more groups used one-way analysis of variance followed by Tukey’s multiple-comparison test. Exact test assignments, significance thresholds, plotted summary statistics and sample numbers are given in the figure legends. For RT–qPCR and dual-luciferase assays, *n* denotes independent biological experiments. For mature-seed plots in Fig. 4a,d and Fig. 5e, each point denotes one seed and the displayed *n* is the number of seeds measured from the representative plant. The seed-level tests describe the displayed representative distributions; the additional independent plants establish reproducibility of phenotype direction but are not pooled as the biological unit in the current panels. The blocked factorial analysis in Fig. 5a and Extended Data Fig. 10 used the two biological sampling blocks as a nuisance factor and controlled the false-discovery rate across all ATH1 probe sets within each contrast.

Sample sizes were chosen with reference to comparable published studies. Samples were not coded, and investigators knew the genotype or group during measurement and analysis.

## Accession numbers

Arabidopsis Genome Initiative identifiers are: *WRKY10/MINI3* (AT1G55600), *IKU1* (AT2G35230), *IKU2* (AT3G19700), *EXPA21* (AT5G39260), *EXPA22* (AT5G39270), *EXPA23* (AT5G39280), *EXPA24* (AT5G39310), *EXPA25* (AT5G39300), *EXPA26* (AT5G39290), *FWA* (AT4G25530), *BAN* (AT1G61720), *RPS5A* (AT3G11940) and *ACT11* (AT3G12110).

## Data availability

The public seed atlas used for Fig. 1a is available from GEO under accession GSE12404^14^. ATAC-seq and microarray data generated in this study have been deposited in GEO under accessions GSE337509, GSE337510 and GSE342468. Processed data underlying Fig. 5a and Extended Data Fig. 10, including the design and contrast matrices, probe-level statistics and software session record, are provided in the accompanying reviewer data and code directory. Plasmids and transgenic lines are available from Jing Li on reasonable request and subject to applicable material-transfer requirements.

## Code availability

Scripts and processed outputs for the blocked factorial analysis in Fig. 5a and Extended Data Fig. 10 are provided in the accompanying reviewer data and code directory. A public repository URL, versioned release and archival identifier will be added before publication.

## Materials availability

Plasmids, plant materials and other reagents generated in this study are available from Jing Li on reasonable request and subject to applicable material-transfer requirements.

## AI-assisted language editing

During preparation of this work, the authors used ChatGPT to improve language and readability. The authors reviewed and edited the resulting text and take full responsibility for the content of the manuscript.

## End Notes

## Acknowledgements

We thank Matt Watson for language proofreading of the manuscript.

## Funding

This work was supported by the National Natural Science Foundation of China (31570322 and 32471538 to J.L.), the Hainan University Startup Fund (RZ2100003224 to J.L.) and the Austrian Science Fund (FWF P36231 to F.B.).

## Author contributions

Conceptualization: J.L., F.B., R.X. and K.Y. Methodology: K.Y., W.B.G., Y.L.T., Y.L., Y.J.Z., X.Y.X., M.M.M., Z.S.W., E.A., H.Y., C.J.X., W.B.X., J.L.Y., Y.Z. and P.T.L. Investigation: K.Y., W.B.G. and J.L. Visualization: K.Y., W.B.G., J.L. and R.X. Funding acquisition: J.L. and F.B. Project administration: J.L. Supervision: J.L., R.X. and F.B. Writing – original draft: J.L., F.B., R.X., K.Y. and W.B.G. Writing – review and editing: J.L., F.B., R.X., K.Y., W.B.G., P.Z. and M.X.S.

## Competing interests

The authors declare no competing interests.

## Additional information

Supplementary Information is available for this paper.

Correspondence should be addressed to Jing Li, Ran Xu or Frédéric Berger. Requests for materials should be addressed to Jing Li.

## Extended Data Figures

**Extended Data Figure 1.**
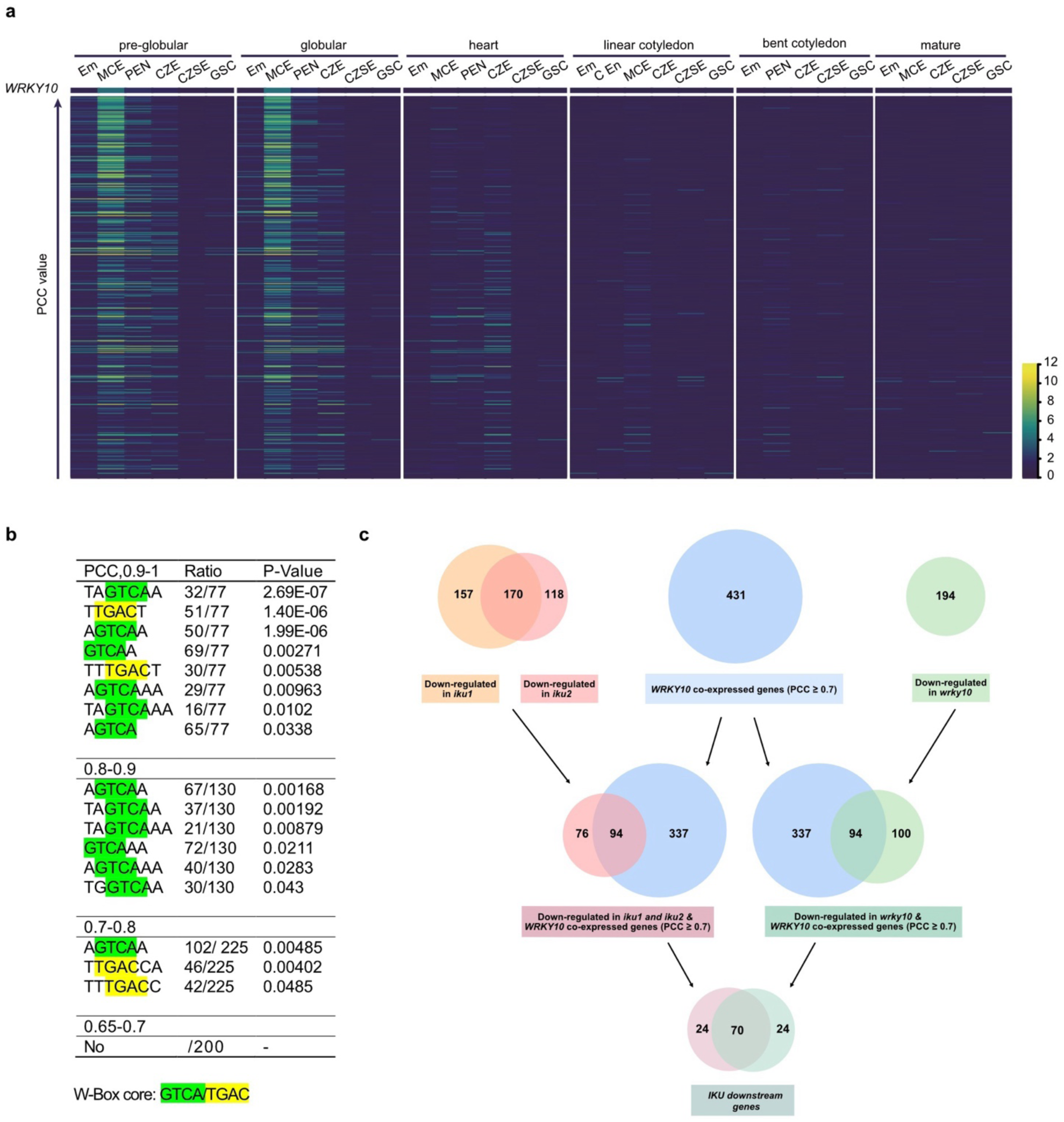
| Definition of the IKU downstream gene set. **a,** Expression profiles of 431 ATH1 probe sets with a Pearson correlation coefficient (PCC) of at least 0.7 with *WRKY10* across seed compartments from the pre-globular to mature stages. Probe sets are ordered by decreasing PCC with *WRKY10*. Colour indicates normalized microarray expression. Em, embryo; MCE, micropylar endosperm; PEN, peripheral endosperm; CZE, chalazal endosperm; CZSE, chalazal seed coat; GSC, general seed coat. **b,** Exact W-box-containing motifs enriched in promoter regions of genes grouped by their PCC with *WRKY10*. Ratios indicate the number of promoters containing each motif over the number of genes in the PCC interval. *P* values report motif enrichment relative to the promoter background used in the analysis. The canonical W-box core is highlighted. **c,** Sequential overlaps among ATH1 probe sets downregulated in *iku1*, *iku2* and *wrky10* and the 431 *WRKY10*-co-expressed probe sets. The final intersection defines 70 IKU downstream genes (*IDGs*).

**Extended Data Figure 2.**
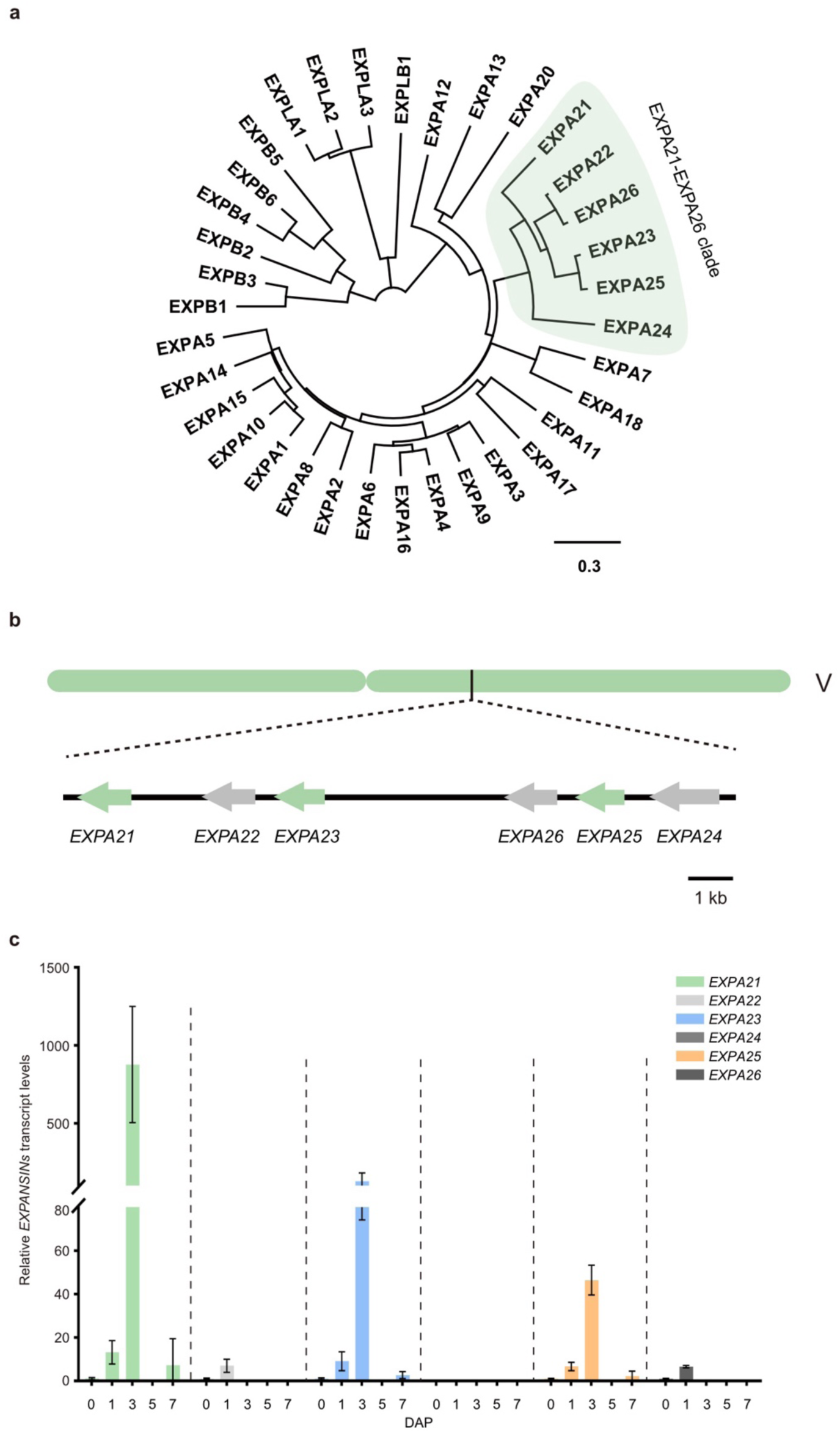
| Phylogeny, genomic organization and developmental expression of the *EXPA21–EXPA26* cluster. **a,** Maximum-likelihood tree of annotated *Arabidopsis* expansins. The clade containing *EXPA21–EXPA26* is shaded. Scale bar, substitutions per site. **b,** Tandem arrangement and transcriptional orientation of *EXPA21– EXPA26* on chromosome 5. Genes analysed in the main text are shown in green. Scale bar, 1 kb. **c,** RT–qPCR analysis of *EXPA21–EXPA26* from unfertilized ovules to 7 days after pollination (DAP). *EXPA21*, *EXPA23* and *EXPA25* are induced after fertilization and reach their highest abundance during early seed development, whereas *EXPA22*, *EXPA24* and *EXPA26* remain low or undetectable. Data are mean ± s.d. of three biological replicates.

**Extended Data Figure 3.**
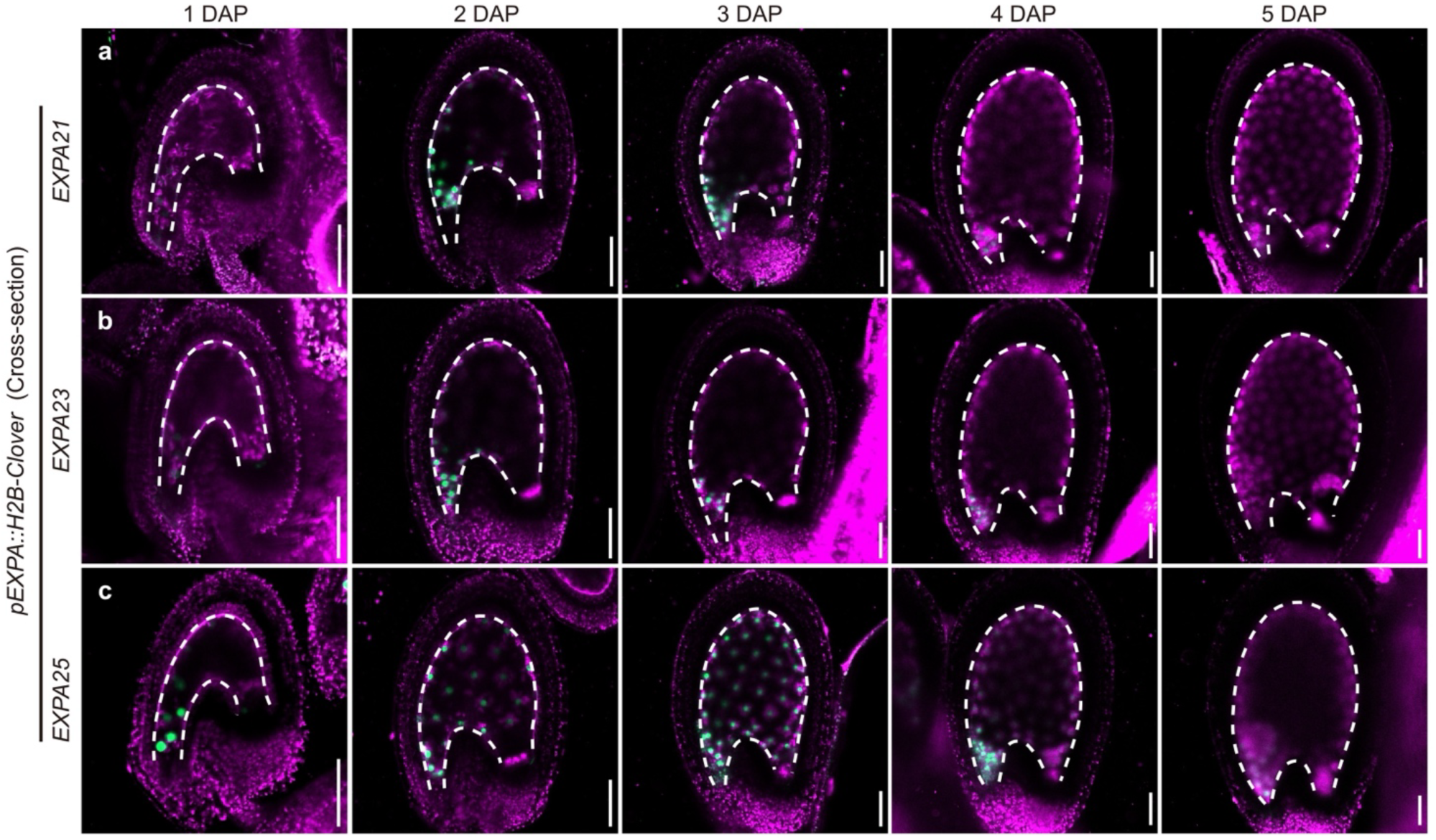
| Spatial and temporal activity of *EXPA21*, *EXPA23* and *EXPA25* promoters. Confocal imaging of *pEXPA21::H2B–Clover* (**a**), *pEXPA23::H2B–Clover* (**b**) and *pEXPA25::H2B–Clover* (**c**) seeds from 1 to 5 DAP. Green, nuclear H2B–Clover signal; magenta, tissue autofluorescence. Dashed lines mark the endosperm-facing inner-integument boundary. Scale bars, 50 μm.

**Extended Data Figure 4.**
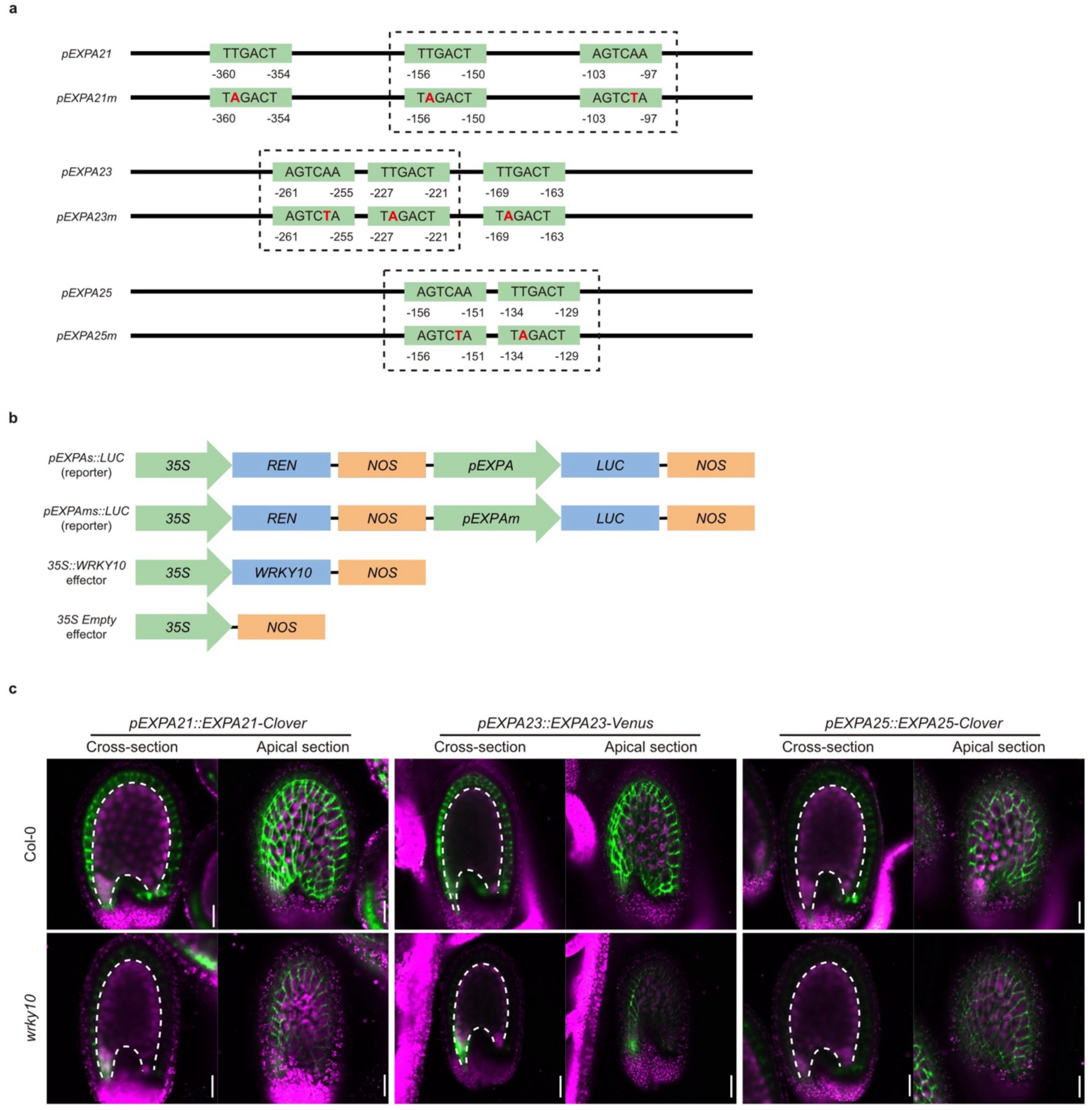
| W-box-dependent activation and WRKY10-dependent accumulation of *EXPA* reporters. **a,** Positions and sequences of W-box motifs in the *EXPA21*, *EXPA23* and *EXPA25* promoter fragments used for electrophoretic mobility-shift and dual-luciferase assays. Mutated bases are shown in red, and dashed boxes indicate the promoter fragments used as binding probes. Coordinates are relative to the translational start site. **b,** Reporter and effector constructs used for dual-luciferase assays. REN, Renilla luciferase; LUC, firefly luciferase; NOS, nopaline synthase terminator. **c,** Cross-sectional and apical views of Col-0 and *wrky10* seeds expressing native-promoter translational reporters for *EXPA21*, *EXPA23* and *EXPA25* at 3 DAP. Green, EXPA–fluorescent-protein signal; magenta, tissue autofluorescence. Dashed lines mark the endosperm-facing inner-integument boundary. Scale bars, 50 μm.

**Extended Data Figure 5.**
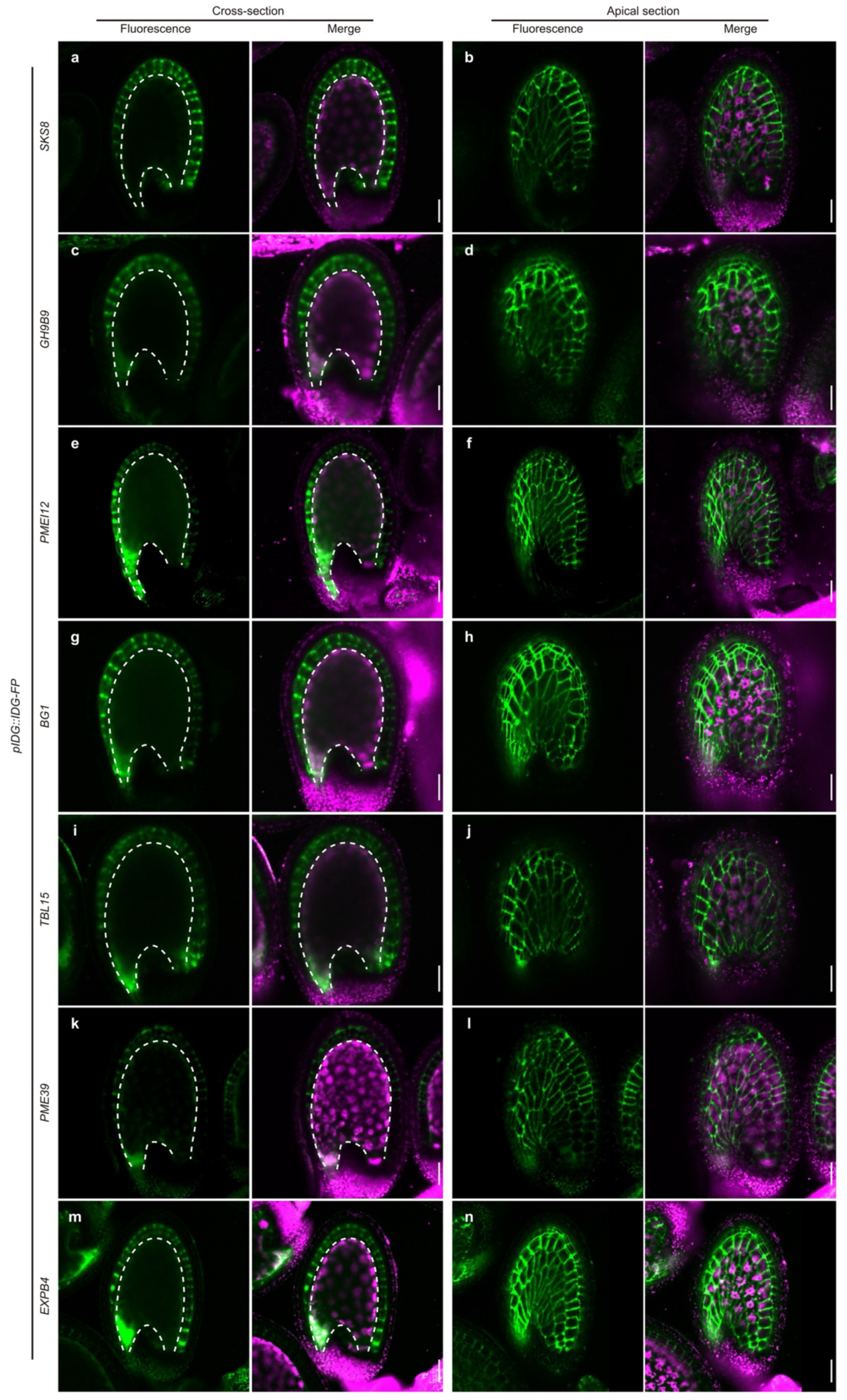
| Additional IKU downstream proteins accumulate in maternal inner-integument walls. Cross-sectional and apical views of 3-DAP seeds expressing native-promoter translational reporters for *SKS8* (**a,b**), *GH9B9* (**c,d**), *PMEI12* (**e,f**), *BG1* (**g,h**), *TBL15* (**i,j**), *PME39* (**k,l**) and *EXPB4* (**m,n**). For each construct, fluorescence and merged views are shown. Green, fusion-protein signal; magenta, tissue autofluorescence. Dashed lines mark the endosperm-facing inner-integument boundary. Scale bars, 50 μm.

**Extended Data Figure 6.**
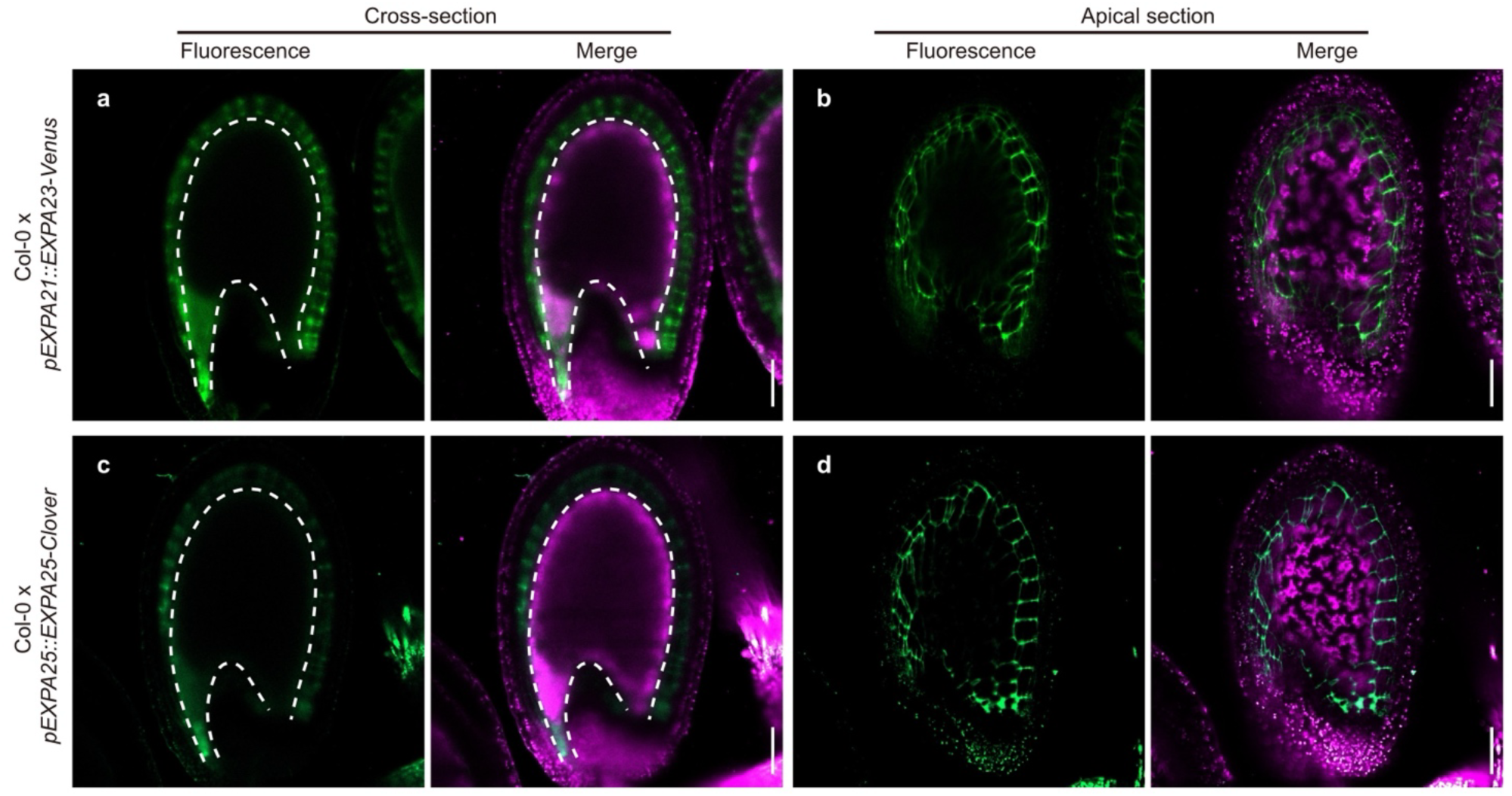
| Paternally inherited *EXPA23* and *EXPA25* reporters accumulate in maternal walls. Cross-sectional and apical views of F1 seeds produced by crossing Col-0 females with pollen donors carrying *pEXPA23::EXPA23–Venus* (**a,b**) or *pEXPA25::EXPA25–Clover* (**c,d**). Fusion proteins expressed from the paternally inherited transgenes accumulate in maternal inner-integument walls. Green, fusion-protein signal; magenta, tissue autofluorescence. Dashed lines mark the endosperm-facing inner-integument boundary. Scale bars, 50 μm.

**Extended Data Figure 7.**
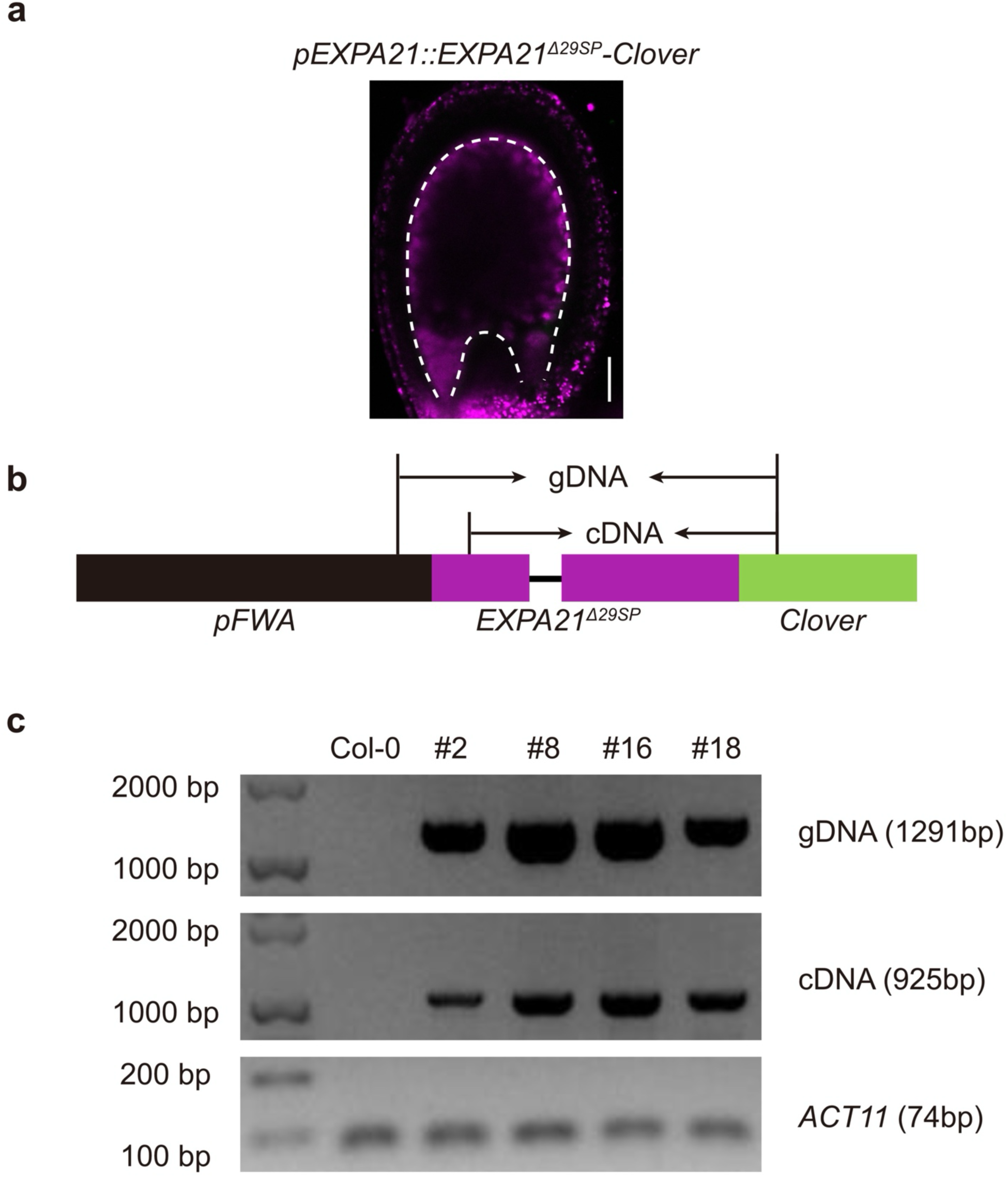
| The EXPA21 signal peptide is required for detectable fusion-protein accumulation. **a,** Confocal image of a seed expressing the native-promoter signal-peptide deletion reporter *pEXPA21::EXPA21^Δ29SP^–Clover*. No Clover fluorescence was detected. Green, Clover signal; magenta, tissue autofluorescence. Dashed lines mark the endosperm-facing inner-integument boundary. Scale bar, 50 μm. **b,** Structure of the independently generated *pFWA::EXPA21^Δ29SP^–Clover* construct and positions of the genomic-DNA and cDNA primer pairs. **c,** Genomic-DNA and cDNA amplification from independent *pFWA::EXPA21^Δ29SP^–Clover* lines. Amplification of the deletion-fusion transcript confirms *FWA*-driven transcription, although no fusion-protein fluorescence was detected. *ACT11* was used as the cDNA control. Product sizes are indicated.

**Extended Data Figure 8.**
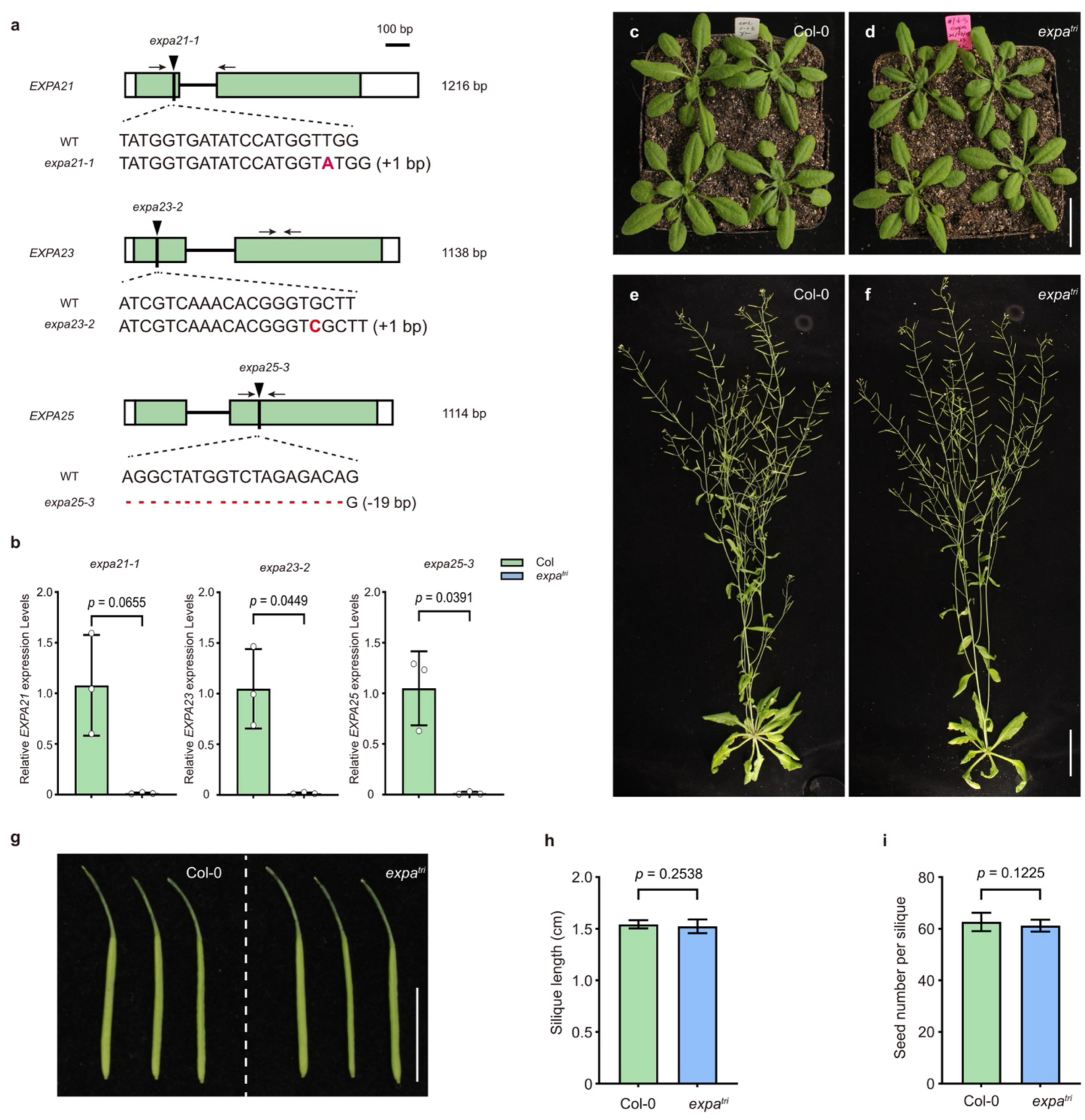
| Generation and vegetative phenotyping of the *expa^tri^* mutant. **a,** Gene structures and edited sequences of *expa21-1*, *expa23-2* and *expa25-3*. Arrowheads mark the edited sites, arrows indicate genotyping-primer positions, and altered bases or deletions are shown in red. **b,** RT–qPCR analysis of *EXPA21*, *EXPA23* and *EXPA25* in Col-0 and the *expa21 expa23 expa25* triple mutant (*expa^tri^*). Data are mean ± s.d. of three biological replicates. Exact *P* values are shown and were calculated using two-tailed Student’s *t*-tests. **c–f,** Col-0 and *expa^tri^*plants at 2 weeks (**c,d**) and 6 weeks (**e,f**) after transplantation. **g,** Representative siliques. **h,i,** Silique length (**h**) and seed number per silique (**i**). Data are mean ± s.d. Exact *P* values are shown and were calculated using two-tailed Student’s *t*-tests. Scale bars are indicated in the images.

**Extended Data Figure 9.**
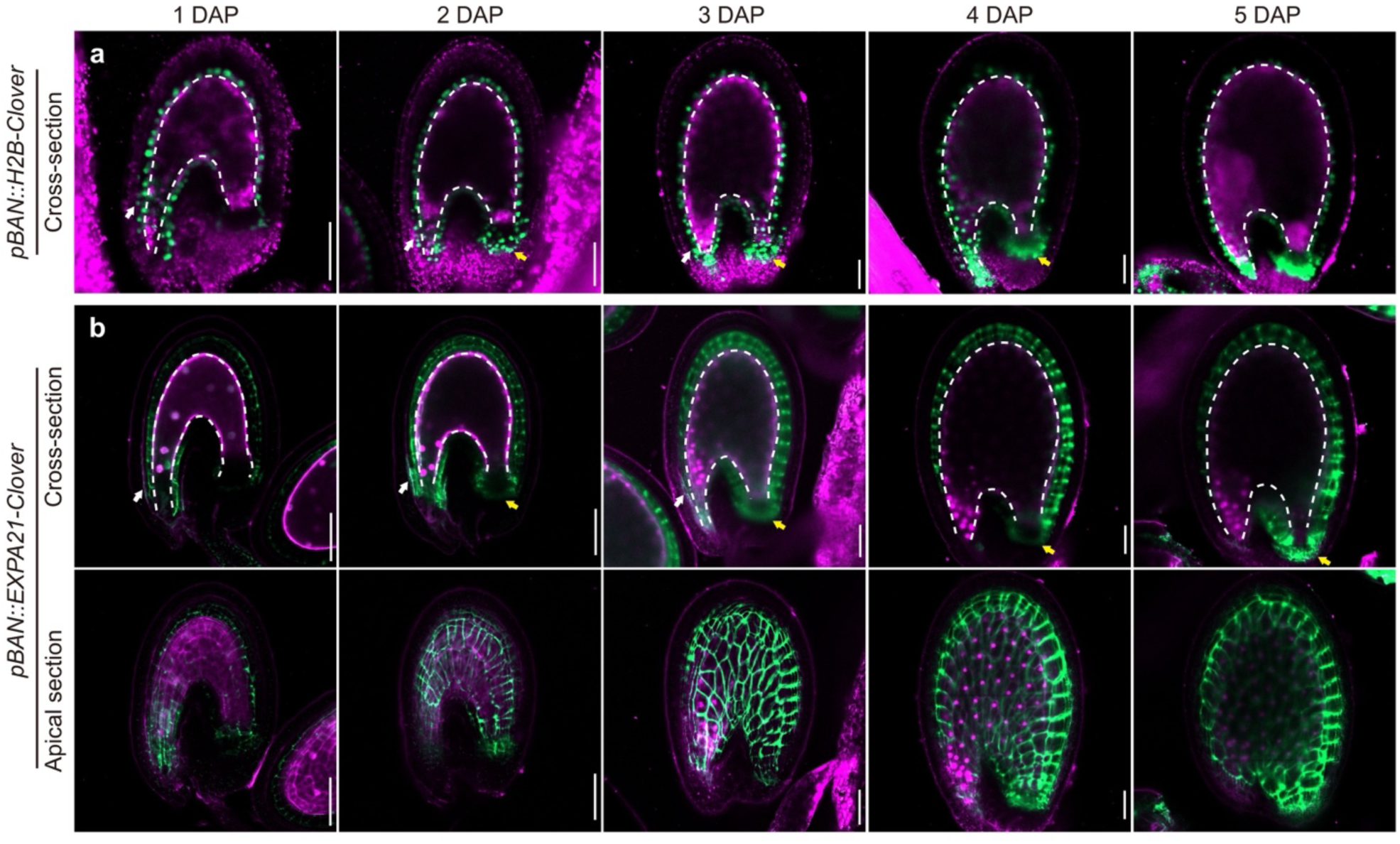
| *BAN*-driven expression and extracellular deployment of EXPA21. **a,** Cross-sectional views of *pBAN::H2B–Clover* seeds from 1 to 5 DAP, showing *BAN* promoter activity in ii1, micropylar ii2 cells and the chalazal pigment strand. **b,** Cross-sectional and apical views of *pBAN::EXPA21– Clover* seeds over the same period. White and yellow arrowheads mark micropylar ii2 and chalazal pigment-strand signal, respectively. Green, Clover signal; magenta, tissue autofluorescence. Dashed lines mark the endosperm-facing inner-integument boundary. Scale bars, 50 μm.

**Extended Data Figure 10.**
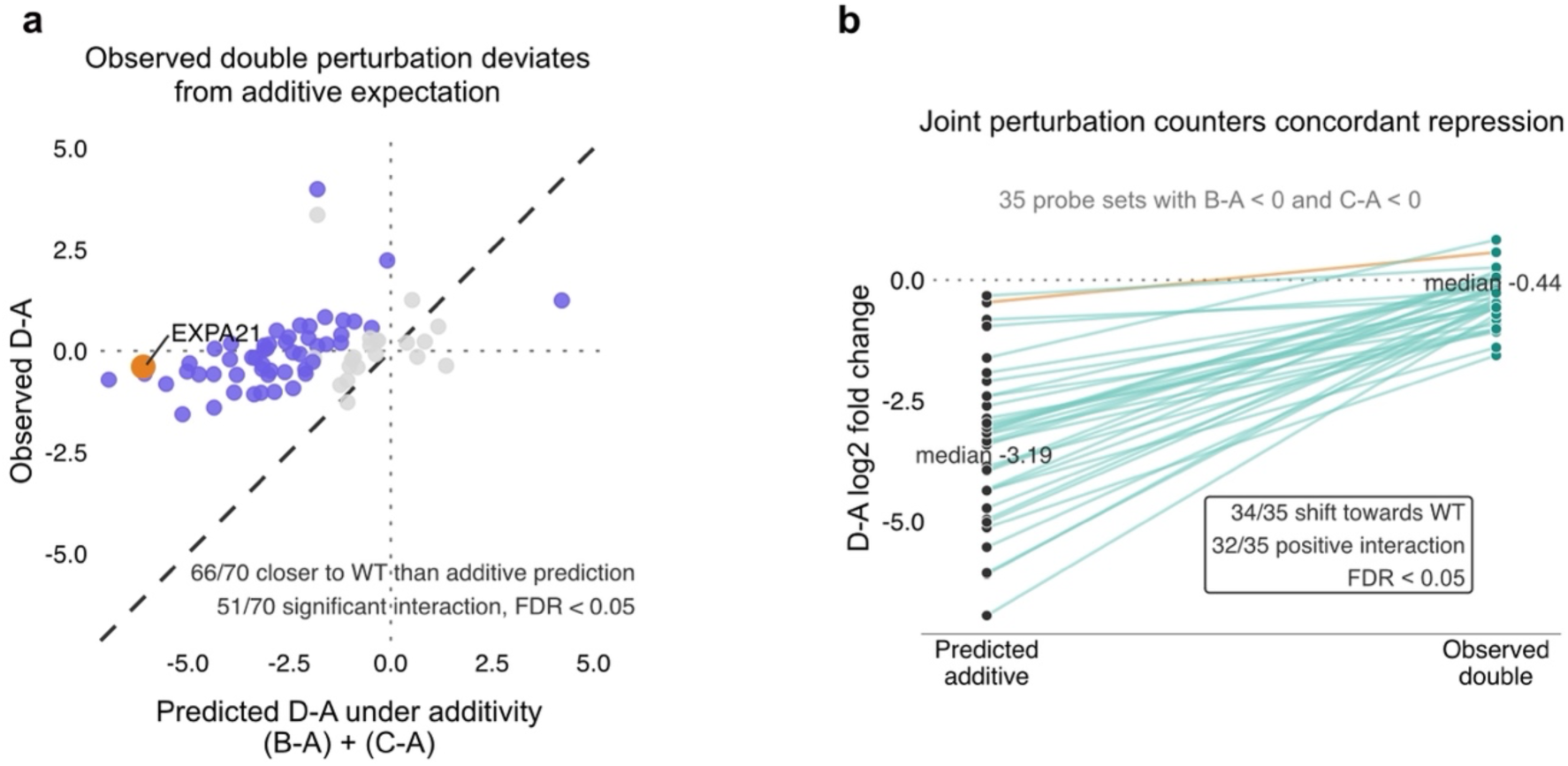
| Non-additive responses of the IKU downstream programme to parental epigenetic perturbations. **a,** Predicted additive response, (B − A) + (C − A), versus the observed double-perturbation response, D − A, for the 70 *IDG*-associated ATH1 probe sets. The four parental conditions are defined in Fig. 5a. The diagonal dashed line indicates the additive expectation, and dotted lines mark zero response. Purple points indicate significant interactions at a false-discovery rate (FDR) < 0.05, grey points indicate the remaining probe sets, and the orange point marks *EXPA21*. Sixty-six of 70 probe sets were closer to wild type than predicted, and 51 had significant interactions. **b,** Predicted and observed responses for the 35 probe sets downregulated in both single perturbations, with B − A < 0 and C − A < 0. Lines connect the two estimates for each probe set. The orange line highlights *EXPA21*. Median values were −3.19 and −0.44 log_2_ units. Thirty-four of 35 probe sets were closer to wild type than predicted, and 32 had positive interactions at FDR < 0.05. A probe set was classified as closer to wild type when |D − A| < |(B − A) + (C − A)|. RMA-normalized data from two biological sampling blocks per cross were analysed using the blocked factorial limma model, with Benjamini–Hochberg adjustment across ATH1 probe sets within each contrast. Complete estimates are provided in Supplementary Table 6.

## Notes

### Competing Interest Statement

The authors have declared no competing interest.

## References

1. Creff, A. et al. Evidence that endosperm turgor pressure both promotes and restricts seed growth and size. Nat. Commun. 14, 67 (2023).

2. Figueiredo, D. D., Batista, R. A., Roszak, P. J., Hennig, L. & Köhler, C. Auxin production in the endosperm drives seed coat development in *Arabidopsis*. eLife 5, e20542 (2016).

3. Lopez-Tello, J. et al. Fetal manipulation of maternal metabolism is a critical function of the imprinted *Igf2* gene. Cell Metab. 35, 1195–1208.e6 (2023).

4. Napso, T., Yong, H. E. J., Lopez-Tello, J. & Sferruzzi-Perri, A. N. The role of placental hormones in mediating maternal adaptations to support pregnancy and lactation. Front. Physiol. 9, 1091 (2018).

5. Greenbaum, S. et al. A spatially resolved timeline of the human maternal–fetal interface. Nature 619, 595–605 (2023).

6. Garcia, D., et al. *Arabidopsis haiku* mutants reveal new controls of seed size by endosperm. Plant Physiol. 131, 1661–1670 (2003).

7. Garcia, D., Fitz Gerald, J. N. & Berger, F. Maternal control of integument cell elongation and zygotic control of endosperm growth are coordinated to determine seed size in *Arabidopsis*. Plant Cell 17, 52–60 (2005).

8. Ingouff, M., Jullien, P. E. & Berger, F. The female gametophyte and the endosperm control cell proliferation and differentiation of the seed coat in *Arabidopsis*. Plant Cell 18, 3491–3501 (2006).

9. Creff, A., Brocard, L. & Ingram, G. A mechanically sensitive cell layer regulates the physical properties of the *Arabidopsis* seed coat. Nat. Commun. 6, 6382 (2015).

10. Bauer, A. et al. Spatiotemporally distinct responses to mechanical forces shape the developing seed of *Arabidopsis*. EMBO J. 43, 2733–2758 (2024).

11. Coen, O. et al. Deposition of a cutin apoplastic barrier separating seed maternal and zygotic tissues. BMC Plant Biol. 19, 304 (2019).

12. Luo, M., Dennis, E. S., Berger, F., Peacock, W. J. & Chaudhury, A. *MINISEED3* (*MINI3*), a WRKY family gene, and *HAIKU2* (*IKU2*), a leucine-rich repeat (LRR) KINASE gene, are regulators of seed size in *Arabidopsis*. Proc. Natl Acad. Sci. USA 102, 17531–17536 (2005).

13. Wang, A. et al. The VQ motif protein IKU1 regulates endosperm growth and seed size in *Arabidopsis*. Plant J. 63, 670–679 (2010).

14. Belmonte, M. F., et al. Comprehensive developmental profiles of gene activity in regions and subregions of the *Arabidopsis* seed. Proc. Natl Acad. Sci. USA 110, E435–E444 (2013).

15. Cosgrove, D. J. Loosening of plant cell walls by expansins. Nature 407, 321–326 (2000).

16. Sampedro, J. & Cosgrove, D. J. The expansin superfamily. Genome Biol. 6, 242 (2005).

17. Kinoshita, T. et al. One-way control of *FWA* imprinting in *Arabidopsis* endosperm by DNA methylation. Science 303, 521–523 (2004).

18. Debeaujon, I. et al. Proanthocyanidin-accumulating cells in *Arabidopsis* testa: regulation of differentiation and role in seed development. Plant Cell 15, 2514–2531 (2003).

19. Coen, O. et al. Developmental patterning of the sub-epidermal integument cell layer in *Arabidopsis* seeds. Development 144, 1490–1497 (2017).

20. Xu, W. et al. Endosperm and nucellus develop antagonistically in *Arabidopsis* seeds. Plant Cell 28, 1343– 1360 (2016).

21. Martin, C. A., Cogdill, K. R., Pusey, A. L. & Gehring, M. A transcriptional atlas of early *Arabidopsis* seed development suggests mechanisms for inter-tissue coordination. Nat. Plants 12, 1133–1149 (2026).

22. Povilus, R. A. & Gehring, M. Maternal-filial transfer structures in endosperm: a nexus of nutritional dynamics and seed development. Curr. Opin. Plant Biol. 65, 102121 (2022).

23. Nguyen, H., Brown, R. C. & Lemmon, B. E. The specialized chalazal endosperm in *Arabidopsis thaliana* and *Lepidium virginicum* (Brassicaceae). Protoplasma 212, 99–110 (2000).

24. Brown, R. C., Lemmon, B. E. & Nguyen, H. Comparative anatomy of the chalazal endosperm cyst in seeds of the Brassicaceae. Bot. J. Linn. Soc. 144, 375–394 (2004).

25. Adams, S., Vinkenoog, R., Spielman, M., Dickinson, H. G. & Scott, R. J. Parent-of-origin effects on seed development in *Arabidopsis thaliana* require DNA methylation. Development 127, 2493–2502 (2000).

26. Luo, M., Bilodeau, P., Dennis, E. S., Peacock, W. J. & Chaudhury, A. Expression and parent-of-origin effects for *FIS2*, *MEA* and *FIE* in the endosperm and embryo of developing *Arabidopsis* seeds. Proc. Natl Acad. Sci. USA 97, 10637–10642 (2000).

27. Moreno-Romero, J., Jiang, H., Santos-González, J. & Köhler, C. Parental epigenetic asymmetry of PRC2-mediated histone modifications in the *Arabidopsis* endosperm. EMBO J. 35, 1298–1311 (2016).

28. Wu, D. et al. Ancestral function but divergent epigenetic regulation of *HAIKU2* reveals routes of seed developmental evolution. Mol. Plant 15, 1575–1589 (2022).

29. Zhang, S., Mohanty, D., Muzaffar, A. & Ni, M. Two MADS-box proteins, AGL9 and AGL15, recruit the FIS–PRC2 complex to trigger the phase transition from endosperm proliferation to embryo development in *Arabidopsis*. Mol. Plant 17, 1110–1128 (2024).

30. Cosgrove, D. J. Plant cell wall loosening by expansins. Annu. Rev. Cell Dev. Biol. 40, 329–352 (2024).

31. Doll, N. M. et al. A two-way molecular dialogue between embryo and endosperm is required for seed development. Science 367, 431–435 (2020).

32. Creff, A. et al. An embryo-derived peptide signal directs endosperm polarity in *Arabidopsis*. Science 393, 1002–1008 (2026).

33. Soh, Y. M., Tiwari, A., Mahendroo, M., Conrad, K. P. & Parry, L. J. Relaxin regulates hyaluronan synthesis and aquaporins in the cervix of late pregnant mice. Endocrinology 153, 6054–6064 (2012).

## Methods References

34. Li, J., Nie, X., Tan, J. L. & Berger, F. Integration of epigenetic and genetic controls of seed size by cytokinin in *Arabidopsis*. Proc. Natl Acad. Sci. USA 110, 15479–15484 (2013).

35. Smyth, D. R., Bowman, J. L. & Meyerowitz, E. M. Early flower development in *Arabidopsis*. Plant Cell 2, 755–767 (1990).

36. Gibson, D. G. et al. Enzymatic assembly of DNA molecules up to several hundred kilobases. Nat. Methods 6, 343–345 (2009).

37. Engler, C., Kandzia, R. & Marillonnet, S. A one pot, one step, precision cloning method with high throughput capability. PLoS ONE 3, e3647 (2008).

38. Xiang, C., Han, P., Lutziger, I., Wang, K. & Oliver, D. J. A mini binary vector series for plant transformation. Plant Mol. Biol. 40, 711–717 (1999).

39. Tsutsui, H. & Higashiyama, T. *pKAMA-ITACHI* vectors for highly efficient CRISPR/Cas9-mediated gene knockout in *Arabidopsis thaliana*. Plant Cell Physiol. 58, 46–56 (2017).

40. Wang, Z.-P. et al. Egg cell-specific promoter-controlled CRISPR/Cas9 efficiently generates homozygous mutants for multiple target genes in *Arabidopsis* in a single generation. Genome Biol. 16, 144 (2015).

41. Xing, H.-L. et al. A CRISPR-Cas9 toolkit for multiplex genome editing in plants. BMC Plant Biol. 14, 327 (2014).

42. Xie, X. et al. CRISPR-GE: a convenient software toolkit for CRISPR-based genome editing. Mol. Plant 10, 1246–1249 (2017).

43. Clough, S. J. & Bent, A. F. Floral dip: a simplified method for *Agrobacterium*-mediated transformation of *Arabidopsis thaliana*. Plant J. 16, 735–743 (1998).

44. Ciolkowski, I., Wanke, D., Birkenbihl, R. P. & Somssich, I. E. Studies on DNA-binding selectivity of WRKY transcription factors lend structural clues into WRKY-domain function. Plant Mol. Biol. 68, 81–92 (2008).

45. Bailey, T. L., Johnson, J., Grant, C. E. & Noble, W. S. The MEME Suite. Nucleic Acids Res. 43, W39–W49 (2015).

46. Teufel, F. et al. SignalP 6.0 predicts all five types of signal peptides using protein language models. Nat. Biotechnol. 40, 1023–1025 (2022).

47. Edgar, R. C. Muscle5: high-accuracy alignment ensembles enable unbiased assessments of sequence homology and phylogeny. Nat. Commun. 13, 6968 (2022).

48. Costa, L. M. et al. Central cell-derived peptides regulate early embryo patterning in flowering plants. Science 344, 168–172 (2014).

49. Yang, K. et al. Two imprinted genes primed by DEMETER in the central cell and activated by WRKY10 in the endosperm. J. Genet. Genomics 51, 855–865 (2024).

50. Hellens, R. P. et al. Transient expression vectors for functional genomics, quantification of promoter activity and RNA silencing in plants. Plant Methods 1, 13 (2005).

51. Kerschen, A., Napoli, C. A., Jorgensen, R. A. & Müller, A. E. Effectiveness of RNA interference in transgenic plants. FEBS Lett. 566, 223–228 (2004).

52. Zhu, T., Liao, K., Zhou, R., Xia, C. & Xie, W. ATAC-seq with unique molecular identifiers improves quantification and footprinting. *Commun*. Biol. 3, 675 (2020).

53. Langmead, B. & Salzberg, S. L. Fast gapped-read alignment with Bowtie 2. Nat. Methods 9, 357–359 (2012).

54. Li, H. et al. The Sequence Alignment/Map format and SAMtools. Bioinformatics 25, 2078–2079 (2009).

55. Zhang, Y. et al. Model-based analysis of ChIP-seq (MACS). Genome Biol. 9, R137 (2008).

56. Schneider, C. A., Rasband, W. S. & Eliceiri, K. W. NIH Image to ImageJ: 25 years of image analysis. Nat. Methods 9, 671–675 (2012).

57. Irizarry, R. A. et al. Exploration, normalization, and summaries of high density oligonucleotide array probe level data. Biostatistics 4, 249–264 (2003).

58. Gautier, L., Cope, L., Bolstad, B. M. & Irizarry, R. A. affy—analysis of Affymetrix GeneChip data at the probe level. Bioinformatics 20, 307–315 (2004).

59. Smyth, G. K. Linear models and empirical Bayes methods for assessing differential expression in microarray experiments. Stat. Appl. Genet. Mol. Biol. 3, Article 3 (2004).

60. Benjamini, Y. & Hochberg, Y. Controlling the false discovery rate: a practical and powerful approach to multiple testing. J. R. Stat. Soc. B 57, 289–300 (1995).

61. Thompson, J. D., Higgins, D. G. & Gibson, T. J. CLUSTAL W: improving the sensitivity of progressive multiple sequence alignment through sequence weighting, position-specific gap penalties and weight-matrix choice. Nucleic Acids Res. 22, 4673–4680 (1994).

62. Kumar, S., Stecher, G., Li, M., Knyaz, C. & Tamura, K. MEGA X: molecular evolutionary genetics analysis across computing platforms. Mol. Biol. Evol. 35, 1547–1549 (2018).

