## Supplementary_Data_and_Code for "Endosperm secretion remodels the maternal seed coat": exact_limma_design_and_contrasts.R

```
sample_info <- data.frame(
  sample_name = c('A1_CC','A2_CC','B1_CM','B2_CM','C1_FC','C2_FC','D1_FM','D2_FM'),
  Block = factor(c('1','2','1','2','1','2','1','2'), levels = c('1','2')),
  Maternal_FIS2 = factor(c('WT','WT','WT','WT','fis2','fis2','fis2','fis2'), levels = c('WT','fis2')),
  Paternal_MET1 = factor(c('WT','WT','met1','met1','WT','WT','met1','met1'), levels = c('WT','met1'))
)
design <- model.matrix(~ Block + Maternal_FIS2 * Paternal_MET1, data = sample_info)
colnames(design) <- make.names(colnames(design))
fit <- lmFit(expr, design)
contrast_matrix <- makeContrasts(
  B_minus_A = Paternal_MET1met1,
  C_minus_A = Maternal_FIS2fis2,
  D_minus_A = Maternal_FIS2fis2 + Paternal_MET1met1 + Maternal_FIS2fis2.Paternal_MET1met1,
  interaction = Maternal_FIS2fis2.Paternal_MET1met1,
  levels = design
)
fit2 <- eBayes(contrasts.fit(fit, contrast_matrix))
```
