## Supplementary_Data_and_Code for "Endosperm secretion remodels the maternal seed coat": IDG70_blocked_summary_statistics.txt

Blocked 2x2 factorial limma model for IDG70 ATH1 probe sets

Model: Expression ~ Block + Maternal_FIS2 + Paternal_MET1 + Maternal_FIS2:Paternal_MET1
FDR values are Benjamini-Hochberg adjusted across all ATH1 probe sets within each contrast.

Number of IDGs with B-A < 0: 69 / 70
Number significantly downregulated in B-A at FDR < 0.05: 62
Median B-A logFC: -2.2789
Number up/down in C-A: up 34, down 36
Number significantly up/down in C-A at FDR < 0.05: up 19, down 20
Median C-A logFC: -0.0436747
Number significantly different in D-A at FDR < 0.05: 15
Median D-A logFC: -0.161425
Number of positive and negative interaction coefficients: positive 63, negative 7
Number of interaction terms significant at FDR < 0.05: 51
Among significant interaction terms, positive 50, negative 1
Observed D-A closer to WT than additive prediction: 66 / 70 (94.3%)
|D-A| < |B-A|: 65 / 70 (92.9%)
|D-A| < |C-A|: 50 / 70 (71.4%)
|D-A| < min(|B-A|, |C-A|): 47 / 70 (67.1%)
