## Supplementary_Data_and_Code for "Endosperm secretion remodels the maternal seed coat": IDG70_both_BA_CA_down_focused_summary.txt

Focused subset: IDG70 probe sets with both B-A < 0 and C-A < 0

Number of such probe sets: 35
Median predicted additive D-A: -3.19476
Median observed D-A: -0.444918
Observed D-A closer to WT than additive prediction: 34 / 35
Significant positive interaction at FDR < 0.05: 32
