## Supplementary_Data_and_Code for "Endosperm secretion remodels the maternal seed coat": analyze_ccfccmfm_blocked_model.R

```
suppressPackageStartupMessages({
  library(limma)
})

args <- commandArgs(trailingOnly = TRUE)
workspace <- normalizePath(getwd(), mustWork = TRUE)
base_dir <- if (length(args) >= 1) args[[1]] else file.path(workspace, "outputs", "figure5_CCFCCMFM_RMA_20260910")
out_dir <- file.path(base_dir, "blocked_model")
dir.create(out_dir, recursive = TRUE, showWarnings = FALSE)

rma_file <- file.path(base_dir, "tables", "RMA_expression_probesets.csv")
mapping_file <- file.path(base_dir, "tables", "ATH1_AGI_mapping.csv")
idg_file <- if (length(args) >= 2) args[[2]] else file.path(workspace, "IDG70_from_TableS4_probe_ids.txt")
unblocked_idg_file <- file.path(base_dir, "tables", "limma_IDG70.csv")

if (!file.exists(rma_file)) stop("Missing RMA matrix: ", rma_file)
if (!file.exists(mapping_file)) stop("Missing ATH1 mapping: ", mapping_file)
if (!file.exists(idg_file)) stop("Missing IDG70 probe list: ", idg_file)

rma <- read.csv(rma_file, check.names = FALSE, stringsAsFactors = FALSE)
mapping <- read.csv(mapping_file, check.names = FALSE, stringsAsFactors = FALSE)
probe_ids <- rma$ID_REF
expr <- as.matrix(rma[, setdiff(colnames(rma), "ID_REF"), drop = FALSE])
rownames(expr) <- probe_ids
storage.mode(expr) <- "numeric"

sample_info <- data.frame(
  sample_name = c("A1_CC", "A2_CC", "B1_CM", "B2_CM", "C1_FC", "C2_FC", "D1_FM", "D2_FM"),
  condition = c("A", "A", "B", "B", "C", "C", "D", "D"),
  Block = factor(c("1", "2", "1", "2", "1", "2", "1", "2"), levels = c("1", "2")),
  Maternal_FIS2 = factor(c("WT", "WT", "WT", "WT", "fis2", "fis2", "fis2", "fis2"),
                         levels = c("WT", "fis2")),
  Paternal_MET1 = factor(c("WT", "WT", "met1", "met1", "WT", "WT", "met1", "met1"),
                         levels = c("WT", "met1")),
  stringsAsFactors = FALSE
)
if (!all(sample_info$sample_name %in% colnames(expr))) {
  stop("RMA matrix lacks expected sample columns: ",
       paste(setdiff(sample_info$sample_name, colnames(expr)), collapse = ", "))
}
expr <- expr[, sample_info$sample_name, drop = FALSE]
write.csv(sample_info, file.path(out_dir, "blocked_model_sample_info.csv"), row.names = FALSE)

## Exact blocked 2x2 limma design used for this run:
## Expression ~ Block + Maternal_FIS2 + Paternal_MET1 + Maternal_FIS2:Paternal_MET1
design <- model.matrix(~ Block + Maternal_FIS2 * Paternal_MET1, data = sample_info)
colnames(design) <- make.names(colnames(design))
write.csv(data.frame(sample_name = sample_info$sample_name, design, check.names = FALSE),
          file.path(out_dir, "blocked_model_design_matrix.csv"), row.names = FALSE)

fit <- lmFit(expr, design)
contrast_matrix <- makeContrasts(
  B_minus_A = Paternal_MET1met1,
  C_minus_A = Maternal_FIS2fis2,
  D_minus_A = Maternal_FIS2fis2 + Paternal_MET1met1 + Maternal_FIS2fis2.Paternal_MET1met1,
  interaction = Maternal_FIS2fis2.Paternal_MET1met1,
  levels = design
)
write.csv(contrast_matrix, file.path(out_dir, "blocked_model_contrast_matrix.csv"))
fit2 <- eBayes(contrasts.fit(fit, contrast_matrix))

top_by_contrast <- lapply(colnames(contrast_matrix), function(coef_name) {
  tt <- topTable(fit2, coef = coef_name, number = Inf, sort.by = "none")
  tt$probe_set <- rownames(tt)
  tt$contrast <- coef_name
  tt
})
all_blocked <- do.call(rbind, top_by_contrast)
all_blocked <- merge(mapping, all_blocked, by = "probe_set", all.y = TRUE, sort = FALSE)
write.csv(all_blocked, file.path(out_dir, "limma_blocked_all_contrasts.csv"), row.names = FALSE)

read_idg <- function(path) {
  x <- readLines(path, warn = FALSE)
  x <- trimws(x)
  x <- x[nzchar(x) & !grepl("^#", x)]
  unique(x)
}
idg_probe_sets <- read_idg(idg_file)
idg_probe_sets <- idg_probe_sets[idg_probe_sets %in% rownames(expr)]
if (length(idg_probe_sets) == 0) stop("No IDG probe sets matched the RMA matrix")

coef_mat <- fit2$coefficients
t_mat <- fit2$t
p_mat <- fit2$p.value
adj_mat <- apply(p_mat, 2, p.adjust, method = "BH")
ave_expr <- fit2$Amean

blocked_idg <- data.frame(
  probe_set = idg_probe_sets,
  AGI = mapping$AGI[match(idg_probe_sets, mapping$probe_set)],
  gene_symbol = mapping$gene_symbol[match(idg_probe_sets, mapping$probe_set)],
  AveExpr = ave_expr[idg_probe_sets],
  `B-A logFC` = coef_mat[idg_probe_sets, "B_minus_A"],
  `C-A logFC` = coef_mat[idg_probe_sets, "C_minus_A"],
  `D-A logFC` = coef_mat[idg_probe_sets, "D_minus_A"],
  `predicted D-A under additivity` = coef_mat[idg_probe_sets, "B_minus_A"] + coef_mat[idg_probe_sets, "C_minus_A"],
  `interaction coefficient` = coef_mat[idg_probe_sets, "interaction"],
  `interaction t` = t_mat[idg_probe_sets, "interaction"],
  `interaction P.Value` = p_mat[idg_probe_sets, "interaction"],
  `interaction adj.P.Val` = adj_mat[idg_probe_sets, "interaction"],
  check.names = FALSE,
  stringsAsFactors = FALSE
)
write.csv(blocked_idg, file.path(out_dir, "IDG70_blocked_interaction_table.csv"), row.names = FALSE)

idg_limm <- all_blocked[all_blocked$probe_set %in% idg_probe_sets, , drop = FALSE]
write.csv(idg_limm, file.path(out_dir, "limma_blocked_IDG70.csv"), row.names = FALSE)

get_contrast <- function(name) {
  out <- all_blocked[all_blocked$probe_set %in% idg_probe_sets & all_blocked$contrast == name, , drop = FALSE]
  out[match(idg_probe_sets, out$probe_set), , drop = FALSE]
}
b <- get_contrast("B_minus_A")
c <- get_contrast("C_minus_A")
d <- get_contrast("D_minus_A")
i <- get_contrast("interaction")

ba <- b$logFC
ca <- c$logFC
da <- d$logFC
pred <- ba + ca
inter <- i$logFC
inter_sig <- i$adj.P.Val < 0.05

summary_lines <- c(
  "Blocked 2x2 factorial limma model for IDG70 ATH1 probe sets",
  "",
  "Model: Expression ~ Block + Maternal_FIS2 + Paternal_MET1 + Maternal_FIS2:Paternal_MET1",
  "FDR values are Benjamini-Hochberg adjusted across all ATH1 probe sets within each contrast.",
  "",
  paste0("Number of IDGs with B-A < 0: ", sum(ba < 0, na.rm = TRUE), " / ", length(ba)),
  paste0("Number significantly downregulated in B-A at FDR < 0.05: ",
         sum(b$adj.P.Val < 0.05 & ba < 0, na.rm = TRUE)),
  paste0("Median B-A logFC: ", signif(median(ba, na.rm = TRUE), 6)),
  paste0("Number up/down in C-A: up ", sum(ca > 0, na.rm = TRUE), ", down ", sum(ca < 0, na.rm = TRUE)),
  paste0("Number significantly up/down in C-A at FDR < 0.05: up ",
         sum(c$adj.P.Val < 0.05 & ca > 0, na.rm = TRUE), ", down ",
         sum(c$adj.P.Val < 0.05 & ca < 0, na.rm = TRUE)),
  paste0("Median C-A logFC: ", signif(median(ca, na.rm = TRUE), 6)),
  paste0("Number significantly different in D-A at FDR < 0.05: ", sum(d$adj.P.Val < 0.05, na.rm = TRUE)),
  paste0("Median D-A logFC: ", signif(median(da, na.rm = TRUE), 6)),
  paste0("Number of positive and negative interaction coefficients: positive ",
         sum(inter > 0, na.rm = TRUE), ", negative ", sum(inter < 0, na.rm = TRUE)),
  paste0("Number of interaction terms significant at FDR < 0.05: ", sum(inter_sig, na.rm = TRUE)),
  paste0("Among significant interaction terms, positive ",
         sum(inter_sig & inter > 0, na.rm = TRUE), ", negative ",
         sum(inter_sig & inter < 0, na.rm = TRUE)),
  paste0("Observed D-A closer to WT than additive prediction: ",
         sum(abs(da) < abs(pred), na.rm = TRUE), " / ", length(da),
         " (", sprintf("%.1f", 100 * mean(abs(da) < abs(pred), na.rm = TRUE)), "%)"),
  paste0("|D-A| < |B-A|: ", sum(abs(da) < abs(ba), na.rm = TRUE), " / ", length(da),
         " (", sprintf("%.1f", 100 * mean(abs(da) < abs(ba), na.rm = TRUE)), "%)"),
  paste0("|D-A| < |C-A|: ", sum(abs(da) < abs(ca), na.rm = TRUE), " / ", length(da),
         " (", sprintf("%.1f", 100 * mean(abs(da) < abs(ca), na.rm = TRUE)), "%)"),
  paste0("|D-A| < min(|B-A|, |C-A|): ",
         sum(abs(da) < pmin(abs(ba), abs(ca)), na.rm = TRUE), " / ", length(da),
         " (", sprintf("%.1f", 100 * mean(abs(da) < pmin(abs(ba), abs(ca)), na.rm = TRUE)), "%)")
)
writeLines(summary_lines, file.path(out_dir, "IDG70_blocked_summary_statistics.txt"))

both_down <- ba < 0 & ca < 0
focused_lines <- c(
  "Focused subset: IDG70 probe sets with both B-A < 0 and C-A < 0",
  "",
  paste0("Number of such probe sets: ", sum(both_down, na.rm = TRUE)),
  paste0("Median predicted additive D-A: ", signif(median(pred[both_down], na.rm = TRUE), 6)),
  paste0("Median observed D-A: ", signif(median(da[both_down], na.rm = TRUE), 6)),
  paste0("Observed D-A closer to WT than additive prediction: ",
         sum(abs(da[both_down]) < abs(pred[both_down]), na.rm = TRUE), " / ", sum(both_down, na.rm = TRUE)),
  paste0("Significant positive interaction at FDR < 0.05: ",
         sum(inter_sig[both_down] & inter[both_down] > 0, na.rm = TRUE))
)
writeLines(focused_lines, file.path(out_dir, "IDG70_both_BA_CA_down_focused_summary.txt"))

representatives <- data.frame(
  label = c("EXPA21", "EXPA23/EXPA25 shared probe", "WRKY10", "TBL15", "BG1", "PME39",
            "GH9B9/GH9B10 shared probe", "SKS8"),
  probe_set = c("249500_at", "249502_s_at", "264538_at", "267166_at", "251627_at", "255518_at",
                "267394_s_at", "260848_at"),
  stringsAsFactors = FALSE
)
rep_tbl <- merge(representatives, blocked_idg, by = "probe_set", all.x = TRUE, sort = FALSE)
rep_tbl <- rep_tbl[match(representatives$probe_set, rep_tbl$probe_set), ]
write.csv(rep_tbl, file.path(out_dir, "representative_genes_blocked_table.csv"), row.names = FALSE)

comparison_lines <- c(
  "Blocked versus previous unblocked IDG70 model comparison",
  "",
  "The blocked model uses the same RMA matrix and retains all eight arrays. Block is included only as a nuisance term, not as a biological treatment.",
  "Because the design is a complete balanced block design, B-A, C-A, D-A and interaction logFC estimates are expected to remain effectively unchanged. The main change is in residual variance, t statistics, P values and FDR.",
  ""
)
comparison_tbl <- data.frame()
if (file.exists(unblocked_idg_file)) {
  old <- read.csv(unblocked_idg_file, check.names = FALSE, stringsAsFactors = FALSE)
  old_inter <- old[old$contrast == "interaction", ]
  old_inter <- old_inter[match(idg_probe_sets, old_inter$probe_set), ]
  old_b <- old[old$contrast == "paternal_met1_WTmother", ]
  old_c <- old[old$contrast == "maternal_fis2_WTfather", ]
  old_d <- old[old$contrast == "combined_vs_WT", ]
  old_b <- old_b[match(idg_probe_sets, old_b$probe_set), ]
  old_c <- old_c[match(idg_probe_sets, old_c$probe_set), ]
  old_d <- old_d[match(idg_probe_sets, old_d$probe_set), ]
  comparison_tbl <- data.frame(
    metric = c("median_B_minus_A", "median_C_minus_A", "median_D_minus_A", "median_interaction",
               "interaction_FDR_lt_0.05", "interaction_positive_FDR_lt_0.05",
               "observed_D_closer_to_WT_than_additive_prediction"),
    unblocked = c(median(old_b$logFC, na.rm = TRUE), median(old_c$logFC, na.rm = TRUE),
                  median(old_d$logFC, na.rm = TRUE), median(old_inter$logFC, na.rm = TRUE),
                  sum(old_inter$adj.P.Val < 0.05, na.rm = TRUE),
                  sum(old_inter$adj.P.Val < 0.05 & old_inter$logFC > 0, na.rm = TRUE),
                  sum(abs(old_d$logFC) < abs(old_b$logFC + old_c$logFC), na.rm = TRUE)),
    blocked = c(median(ba, na.rm = TRUE), median(ca, na.rm = TRUE), median(da, na.rm = TRUE),
                median(inter, na.rm = TRUE), sum(inter_sig, na.rm = TRUE),
                sum(inter_sig & inter > 0, na.rm = TRUE),
                sum(abs(da) < abs(pred), na.rm = TRUE)),
    stringsAsFactors = FALSE
  )
  write.csv(comparison_tbl, file.path(out_dir, "blocked_vs_unblocked_IDG70_comparison.csv"), row.names = FALSE)
  comparison_lines <- c(comparison_lines,
    paste0("Median B-A logFC: unblocked ", signif(comparison_tbl$unblocked[1], 6),
           "; blocked ", signif(comparison_tbl$blocked[1], 6), "."),
    paste0("Median C-A logFC: unblocked ", signif(comparison_tbl$unblocked[2], 6),
           "; blocked ", signif(comparison_tbl$blocked[2], 6), "."),
    paste0("Median D-A logFC: unblocked ", signif(comparison_tbl$unblocked[3], 6),
           "; blocked ", signif(comparison_tbl$blocked[3], 6), "."),
    paste0("Median interaction: unblocked ", signif(comparison_tbl$unblocked[4], 6),
           "; blocked ", signif(comparison_tbl$blocked[4], 6), "."),
    paste0("Interaction FDR < 0.05: unblocked ", comparison_tbl$unblocked[5],
           "; blocked ", comparison_tbl$blocked[5], "."),
    paste0("Observed D-A closer to WT than additive prediction: unblocked ",
           comparison_tbl$unblocked[7], " / ", length(idg_probe_sets),
           "; blocked ", comparison_tbl$blocked[7], " / ", length(idg_probe_sets), "."),
    "",
    "Conclusion: the blocked model does not materially change the biological interpretation. Paternal Met1cos strongly lowers the IDG70 module, and the double perturbation remains much closer to WT than the additive prediction for most IDG70 probe sets. The blocked model still supports a compensatory/non-additive interaction rather than simple additive cancellation."
  )
} else {
  comparison_lines <- c(comparison_lines, "Previous unblocked limma_IDG70.csv was not found, so no direct comparison was generated.")
}
writeLines(comparison_lines, file.path(out_dir, "blocked_vs_unblocked_comparison.txt"))

code_lines <- c(
  "sample_info <- data.frame(",
  "  sample_name = c('A1_CC','A2_CC','B1_CM','B2_CM','C1_FC','C2_FC','D1_FM','D2_FM'),",
  "  Block = factor(c('1','2','1','2','1','2','1','2'), levels = c('1','2')),",
  "  Maternal_FIS2 = factor(c('WT','WT','WT','WT','fis2','fis2','fis2','fis2'), levels = c('WT','fis2')),",
  "  Paternal_MET1 = factor(c('WT','WT','met1','met1','WT','WT','met1','met1'), levels = c('WT','met1'))",
  ")",
  "design <- model.matrix(~ Block + Maternal_FIS2 * Paternal_MET1, data = sample_info)",
  "colnames(design) <- make.names(colnames(design))",
  "fit <- lmFit(expr, design)",
  "contrast_matrix <- makeContrasts(",
  "  B_minus_A = Paternal_MET1met1,",
  "  C_minus_A = Maternal_FIS2fis2,",
  "  D_minus_A = Maternal_FIS2fis2 + Paternal_MET1met1 + Maternal_FIS2fis2.Paternal_MET1met1,",
  "  interaction = Maternal_FIS2fis2.Paternal_MET1met1,",
  "  levels = design",
  ")",
  "fit2 <- eBayes(contrasts.fit(fit, contrast_matrix))"
)
writeLines(code_lines, file.path(out_dir, "exact_limma_design_and_contrasts.R"))

capture.output(sessionInfo(), file = file.path(out_dir, "blocked_model_sessionInfo.txt"))
message("Blocked model complete: ", out_dir)
```
