## Supplementary_Data_and_Code for "Endosperm secretion remodels the maternal seed coat": README.md

# Supporting analysis data and code

This archive accompanies “Endosperm secretion remodels the maternal seed coat”. It contains the recorded inputs, scripts and results for the sampling-block-adjusted analysis underlying Figure 5a, Extended Data Figure 10 and Supplementary Table 6. No new statistical analysis was performed when preparing this archive.

The enclosing directory is named `reviewer_data_and_code` to match the accompanying-directory terminology in the author-submitted manuscript. Its scientific analysis files are supplied here as public supporting material. It contains no reviewer access credentials or private correspondence.

The authoritative analysis script is `figure5/scripts/analyze_ccfccmfm_blocked_model.R`. The historical unblocked script is not included. Recorded results are retained in `figure5/processed`. The four contrasts in `limma_blocked_IDG70.csv` are the numerical source for Supplementary Table 6.

To run the recorded analysis in a separate working copy, with R and limma installed, from this directory:

```bash
Rscript figure5/scripts/analyze_ccfccmfm_blocked_model.R figure5/input_base figure5/inputs/IDG70_from_TableS4_probe_ids.txt
```

The script writes to `figure5/input_base/blocked_model`, leaving the archived `figure5/processed` results intact. Original sequencing and array data are identified by the GEO accessions in the manuscript; this archive does not replace those deposits. It is not a complete collection of the original microscopy or phenotype measurements.
